# The Hrq1 helicase stimulates Pso2 translesion nuclease activity to promote DNA inter-strand crosslink repair

**DOI:** 10.1101/773267

**Authors:** Cody M. Rogers, Chun-Ying Lee, Samuel Parkins, Nicholas J. Buehler, Sabine Wenzel, Francisco Martínez-Márquez, Yuichiro Takagi, Sua Myong, Matthew L. Bochman

## Abstract

DNA inter-strand crosslink (ICL) repair requires a complicated network of DNA damage response pathways. Removal of these lesions is vital as they are physical barriers to essential DNA processes that require the separation of duplex DNA, such as replication and transcription. The Fanconi anemia (FA) pathway is the principle mechanism for ICL repair in metazoans and is coupled to replication. In *Saccharomyces cerevisiae*, a degenerate FA pathway is present, but ICLs are predominantly repaired by a pathway involving the Pso2 nuclease that is hypothesized to digest through the lesion to provide access for translesion polymerases. However, Pso2 lacks translesion nuclease activity *in vitro*, and mechanistic details of this pathway are lacking, especially relative to FA. We recently identified the Hrq1 helicase, a homolog of the disease-linked RECQL4, as a novel component of Pso2- mediated ICL repair. Here, we show that Hrq1 stimulates the Pso2 nuclease in a mechanism that requires Hrq1 catalytic activity. Importantly, Hrq1 also stimulates Pso2 translesion nuclease activity through a site-specific ICL *in vitro*. Stimulation of Pso2 nuclease activity is specific to eukaryotic RecQ4 subfamily helicases, and Hrq1 likely interacts with Pso2 through their N-terminal domains. These results advance our understanding of FA-independent ICL repair and establish a role for the RecQ4 helicases in the repair of these dangerous lesions.

## Introduction

DNA inter-strand crosslinks (ICLs) are covalent linkages between complementary DNA strands, and they act as physical barriers to essential DNA transactions like replication and transcription (1). Repair of these lesions is vital for cell survival, and 20-40 lesions are lethal to repair-deficient mammalian cells (2). For this reason, DNA damaging agents that cause ICLs are common chemotherapeutics, and upregulation of pathways that repair these lesions is a known source of chemotherapeutic resistance (1). To date, the most thoroughly studied ICL repair mechanism involves the Fanconi anemia (FA) pathway in which over 20 proteins are involved (3). The main mechanism for FA-dependent ICL repair is coupled to DNA replication. Briefly, the replisome stalls 20-40 nts from the ICL, which results in uncoupling of the MCM2-7 replicative helicase and DNA synthesis to within 1 nt of the lesion (4). Unhooking of the lesion is accomplished by a suite of nucleases that act in a context-specific manner (reviewed in (5)). This results in a DNA double-strand break (DSB), which is repaired by homologous recombination (HR). Subsequently, translesion polymerases replicate past the remaining ICL adduct, and nucleotide excision repair (NER) factors remove the remaining adducted nucleotide.

However, it has also been shown that a large number of ICLs can be bypassed by an intact replisome in a traverse model in a FA-dependent manner (6). Variations of this mechanism are dependent on the context in which the ICL is identified, and the FA pathway only accounts for ICL repair during S-phase. Importantly, there are numerous ICL repair factors that do not fall within the FA complementation group, including proteins in the SAN1/SETX pathway (7), base excision repair-associated ICL repair (8,9), the repair of acetaldehyde-derived ICLs (10), and the SNM1/Pso2 family nucleases (11). Taken together, ICL repair requires the complex coordination of multiple pathways that depend on the context of the lesion.

Of the three SNM1 proteins in humans (SNM1A, SNM1B (Apollo), and SNM1C (Artemis)), SNM1A is the most directly linked to ICL repair (11), though SNM1B has a role in ICL repair that is independent of SNM1A (12). The model for SNM1A in ICL repair starts with FAN1 and NER factors such as XPF-ERCC1 using their endonuclease activity to create a single-stranded (ss)DNA nick on either the 5′ side of the lesion or on both sides, though other nucleases have been implicated in this process (reviewed in (5)). SNM1A uses its 5′ → 3′ exonuclease activity to digest from the incision through the ICL, facilitating gap fill-in by translesion synthesis (TLS) DNA polymerases (13). While SNM1A appears to play an important role in ICL repair in vertebrates, the FA pathway is the dominant mechanism for ICL repair. However, the *Saccharomyces cerevisiae* homolog of SNM1A, Pso2, is involved in the predominant pathway for ICL repair in yeast (14,15). Indeed, human SNM1A is able to suppress the sensitivity of *pso2Δ* cells to ICL damage (16). Similar to its human counterpart, Pso2 possesses 5′ → 3′ exonuclease activity (17), and is reported to have structure-specific endonuclease activity when used at high concentrations *in vitro* (18). However, Pso2 lacks the ability to digest through an ICL *in vitro*, making the mechanism ICL repair via the Pso2 pathway unclear. We recently identified the *S. cerevisiae* RECQL4 homolog, the Hrq1 helicase, as an additional component of the Pso2-dependent ICL repair pathway (14) and are seeking to further define its role.

The RecQ family helicases are conserved mediators of genome stability, with five family members encoded by the human genome (reviewed in (19)). Mutations in three of the human RecQ helicases (BLM, WRN, and RECQL4) are directly linked to diseases that clinically overlap in their predisposition to cancer and premature aging phenotypes. The involvement of RECQL4 in ICL repair is unclear (20), largely due to technical challenges associated with RECQL4 analysis. Since the identification of RecQ4 helicases in various fungal and plants species (21), new homologs have been identified in bacteria and archaea (22,23), making RECQL4 the only RecQ subfamily helicase conserved in all three domains of life. Recent work on the RECQL4 homologs in *Arabidopsis thaliana* (24) and *S. cerevisiae* (14,25), both called Hrq1, demonstrates that they are involved in ICL repair, similar to RECQL4 (26). Furthermore, Hrq1 appears to be a *bona fide* RECQL4 homolog *in vitro* and *in vivo*, making it a good model for RECQL4-mediated DNA repair (14,25,27). Hrq1 is currently the only known protein to work with Pso2 at the post-incision step of ICL repair, but its mechanism of action is unclear.

Here, we provide further evidence that Hrq1 functions alongside Pso2 to repair ICL lesions that vary in their DNA sequence preference and effect on DNA structure. *In vitro*, Hrq1 stimulated Pso2 nuclease activity in a reaction that requires Hrq1 catalytic activity, and this phenomenon was specific to RecQ4 subfamily helicases. Importantly, we also found that Pso2 stalling at a site-specific ICL can be overcome in the presence of Hrq1. Finally, we demonstrate that the Pso2 N-terminus is an autoinhibitory domain that may act as the interaction platform for Hrq1-mediated nuclease stimulation. These data support the direct role of RecQ4 family helicases in ICL processing and provide mechanistic insight into the Pso2-dependent ICL repair pathway.

## Results

### Hrq1 and Pso2 repair a variety of ICLs

All ICLs are covalent linkages between complementary strands of DNA, but ICL-inducing agents vary in DNA sequence preference and how they affect DNA structure (reviewed in (28)). Indeed, ICL repair pathway utilization in mammals varies depending on the types of crosslinkers being used. For example, highly DNA-distorting lesions like cisplatin and nitrogen mustard ICLs are repaired via the canonical FA pathway (4), whereas psoralen- and abasic site-induced ICLs are preferentially unhooked via the NEIL3 DNA glycosylase in a FA-independent manner (29). To determine if the Pso2-dependent ICL repair pathway in *S. cerevisiae* is dependent on the type of ICL formed, we tested the sensitivity of *pso2* mutants to several ICL-inducing agents. First, we examined the sensitivity of *pso2Δ* cells to mitomycin C (MMC), diepoxybutane (DEB), and 8-methoxypsoralen (8-MOP) + UVA. As diagrammed in Figure S1A, MMC does not distort the DNA backbone, DEB bends the DNA, and 8-MOP + UVA leads to ∼25° unwinding of the DNA around the lesion. Further, all three ICL inducing agents target different DNA sequences. Regardless of the type of ICL formed, deletion of *PSO2* severely sensitized cells to each type of ICL (Fig. 1 and S1). Similar to cells lacking *PSO2* and previously reported results (14,25), mutation of *HRQ1* also rendered cells sensitive to the various ICL-inducing agents (Fig. 1 and S1), though Hrq1 does not appear to be as important in ICL repair as Pso2 because *hrq1Δ* cells were less sensitive to ICL damage than *pso2Δ* cells. Importantly, the deletion of both *HRQ1* and *PSO2* phenocopied the *pso2Δ* levels of MMC (Fig. 1A) and DEB sensitivity (Fig. 1B), consistent with Hrq1 functioning in the Pso2- dependent pathway. The observed sensitivity of these mutants is specific to ICL damage as neither *pso2Δ* nor *hrq1Δ* cells were sensitive to the DNA alkylating agent methyl methanesulfonate (MMS) (30) (Fig. S1A).

**Figure 1.**
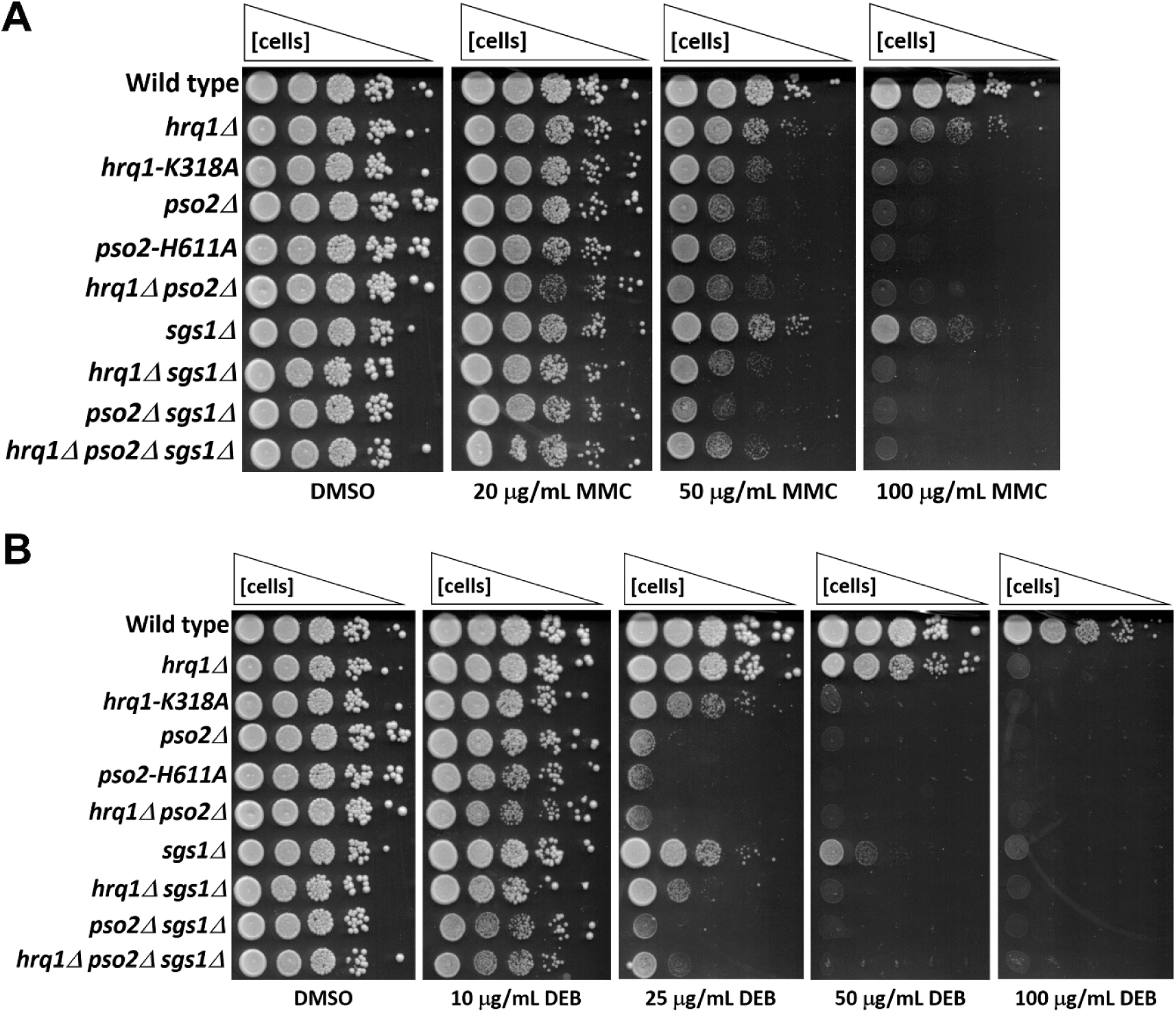
Hrq1 and Pso2 participate in the same ICL repair pathway for MMC and DEB lesions. **A**. Mutation of *HRQ1* is epistatic to *pso2* for MMC sensitivity. Saturated overnight cultures of strains with the genotypes indicated on the left were diluted to OD_660_ = 1.0 and then further serially diluted 10-fold to 10^−4^. Equal volumes of each dilution were then spotted onto rich medium containing the solvent control (DMSO) or rich medium supplemented with the indicated concentration of MMC. **B**. Mutation of *HRQ1* is epistatic to *pso2* for DEB sensitivity. The assay was performed as described above, and the DMSO control plate is shown again for ease of comparison. Mutation of *SGS1* is not epistatic with either *hrq1* or *pso2* in these assays. These results are representative of ≥ 3 independent experiments.

Hrq1 helicase activity is required for the repair of MMC ICLs (25), so we also tested the sensitivity of cells encoding *hrq1-K318A*, a helicase-inactive mutant of Hrq1, to DEB and 8-MOP + UVA. We found that the *hrq1-K318A* mutant is more sensitive to ICLs compared to the *hrq1Δ* strain (Fig. 1 and S1). These results suggest that Hrq1-K318A is recruited to the lesion but is perhaps blocking Pso2 or other redundant repair pathways from accessing the site of damage. Because the *hrq1*-*K318A* strain is slightly less sensitive than *pso2Δ*, however, some amount of Pso2 or a compensatory pathway likely still has access to repair the lesion. This contrasts with the *pso2*-*H611A* nuclease-inactive mutant, which phenocopies *pso2Δ*. These results suggest that Hrq1 may be directly recruited to DNA ICLs and that it may work with Pso2 to repair these lesions.

### Hrq1 stimulates Pso2 nuclease activity

Our genetic analyses suggest that Hrq1 may have a direct role in ICL processing alongside Pso2. Indeed, there is a rich literature of RecQ family helicases and nucleases working in tandem (for instance, see: (31,32)). To investigate this further, we purified recombinant Pso2 and tested it for nuclease activity on blunt dsDNA. Similar to previous work with Pso2 (18), we observed 5′ phosphate-dependent 5′ → 3′ exonuclease activity (Fig. 2A). Here, only the radiolabelled oligonucleotide was 5′-phosphorylated to select for digestion of one strand (Fig. S2A). In the presence of Hrq1, Pso2 nuclease activity increased in a concentration dependent manner up to a nearly threefold stimulation (Fig. 2B). At 200 nM, Pso2 digested ∼80% of the full-length substrate, while 50 nM Pso2 was able to digest nearly as much DNA upon the addition of 150 nM Hrq1.

**Figure 2.**
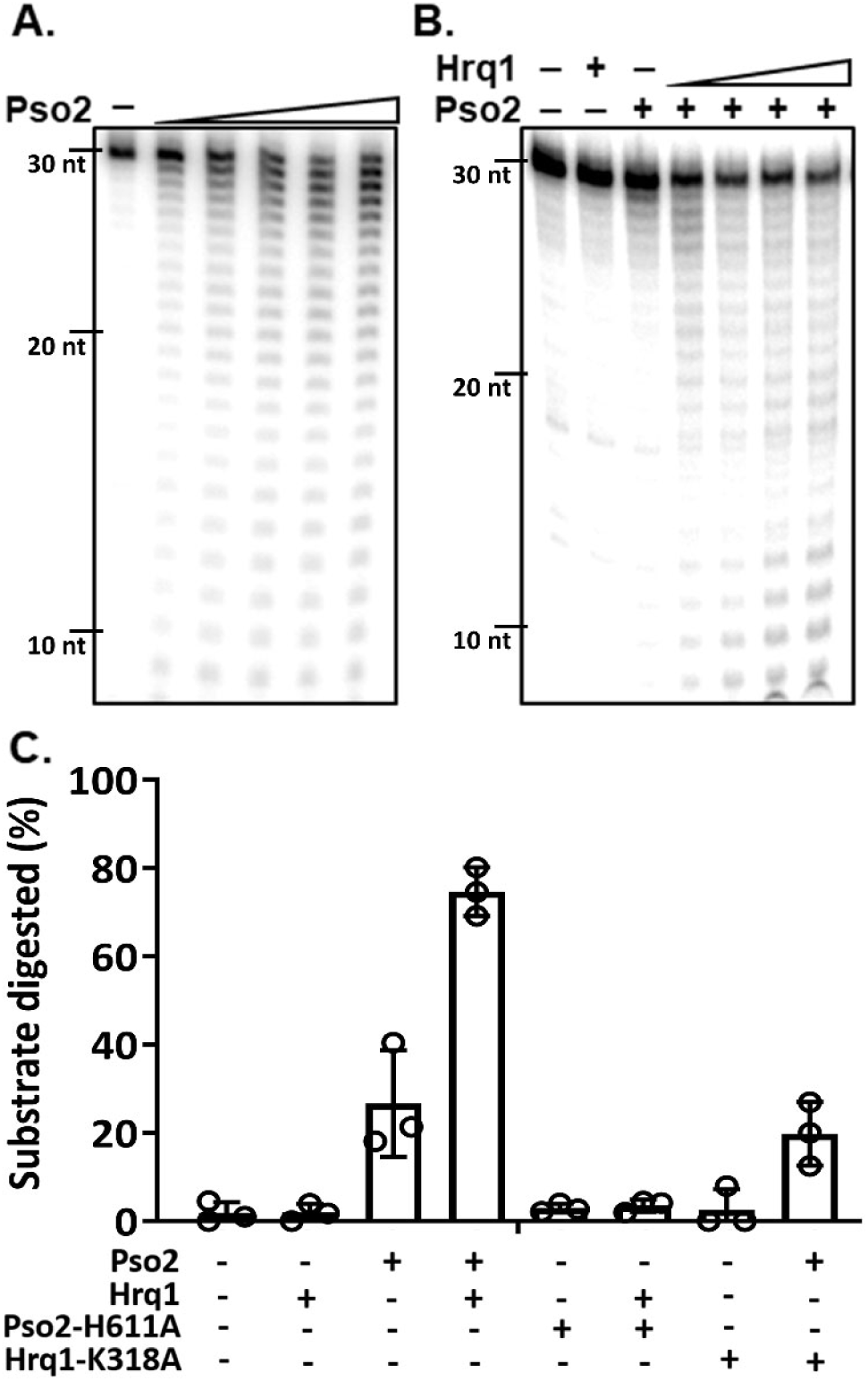
Hrq1 stimulates Pso2 nuclease activity. **A**. Concentration-dependent nuclease activity. Recombinant Pso2 (20-200 nM) was incubated with dsDNA for 30 min, and the nuclease products were separated on a denaturing gel. **B**. Hrq1-dependent stimulation of Pso2 nuclease activity. Denaturing gel showing the radiolabelled dsDNA substrate incubated alone, with 50 nM Pso2, 150 nM Hrq1, or 50 nM Pso2 and 20-150 nM Hrq1. **C**. Hrq1 requires catalytic activity to stimulate Pso2 nuclease activity. Quantification of the nuclease activity of Pso2 alone and in the presence of Hrq1 or the inactive Hrq1-K318A mutant. Nuclease-inactive Pso2-H611A was used as a control. The graphed bars are the averages of three independent experiments (individual data points shown as open circles), and the error bars are the standard deviation (S.D.).

To verify that our observed nuclease activity was due to Pso2 and not a contaminant from *E. coli*, we purified the nuclease-null Pso2-H611A mutant and tested its nuclease activity. Importantly, Pso2-H611A showed no nuclease activity above background, and no stimulation of the mutant was observed in the presence of Hrq1 (Fig. 2C and S2B). Because Hrq1 helicase activity was required for ICL repair *in vivo* (Fig. 1 and S1), we also assayed the catalytically inactive Hrq1-K318A for stimulation of Pso2 nuclease activity *in vitro*. Hrq1-K318A was unable to increase Pso2 nuclease activity, suggesting that Hrq1 catalytic activity is essential to stimulate Pso2 (Fig. S2B). Taken together, the data indicate that Hrq1 stimulates the exonuclease activity of Pso2.

### Hrq1 promotes Pso2 digestion through an ICL

It has been reported that Pso2 lacks translesional nuclease activity *in vitro*, being unable to degrade DNA past a site-specific ICL (18). However, the human homolog of Pso2, SNM1A, does display *in vitro* translesion exonuclease activity (13). Because Hrq1 can stimulate Pso2 nuclease activity on undamaged DNA, we hypothesized that Hrq1 may also stimulate its translesional nuclease activity across an ICL. Thus, we next performed nuclease assays with a dsDNA substrate containing an ICL 7-nt from the 5′ end of the phosphorylated (*i*.*e*., the digested) strand. To make this substrate, we used an established protocol to form ICLs from abasic sites *in vitro* (33). These ICLs are site specific and produced at high yields (up to ∼70% crosslinks), which is advantageous relative to the difficult-to-make drug-based ICLs (28). Semlow *et al*. show that the repair of ICLs from abasic sites proceeds via the same mechanism that repairs psoralen ICLs (29), suggesting that this type of lesion is functionally equivalent to ICLs induced by small molecules.

Due to difficulties in reversing the crosslink in our substrate for denaturing PAGE analysis, gel-based assays of Pso2 translesion nuclease activity were inconclusive (data not shown). To investigate this in a more definitive manner, we instead utilized single-molecule Förster resonance energy transfer (smFRET) to measure Pso2 nuclease activity. To observe nuclease activity, we prepared a DNA substrate that contains Cy3, Cy5, and biotin in the undigested strand, which was annealed to an unlabelled but 5’-phosphorylated strand to be digested by Pso2 (Fig. 3A). Because the FRET efficiency reports on the distance between the donor and acceptor, the FRET signals of dsDNA and ssDNA can be distinguished as low and high FRET, respectively (34). Therefore, the changes in FRET signal as ssDNA is generated by Pso2 digestion of the unlabelled strand can be used to measure nuclease activity. Thus, we expected to see no change in FRET in the initial phase of Pso2 loading, followed by FRET increase induced by Pso2 digestion and concomitant generation of ssDNA and subsequent high FRET state resulting from the completion of the digestion (Fig. 3A).

**Figure 3.**
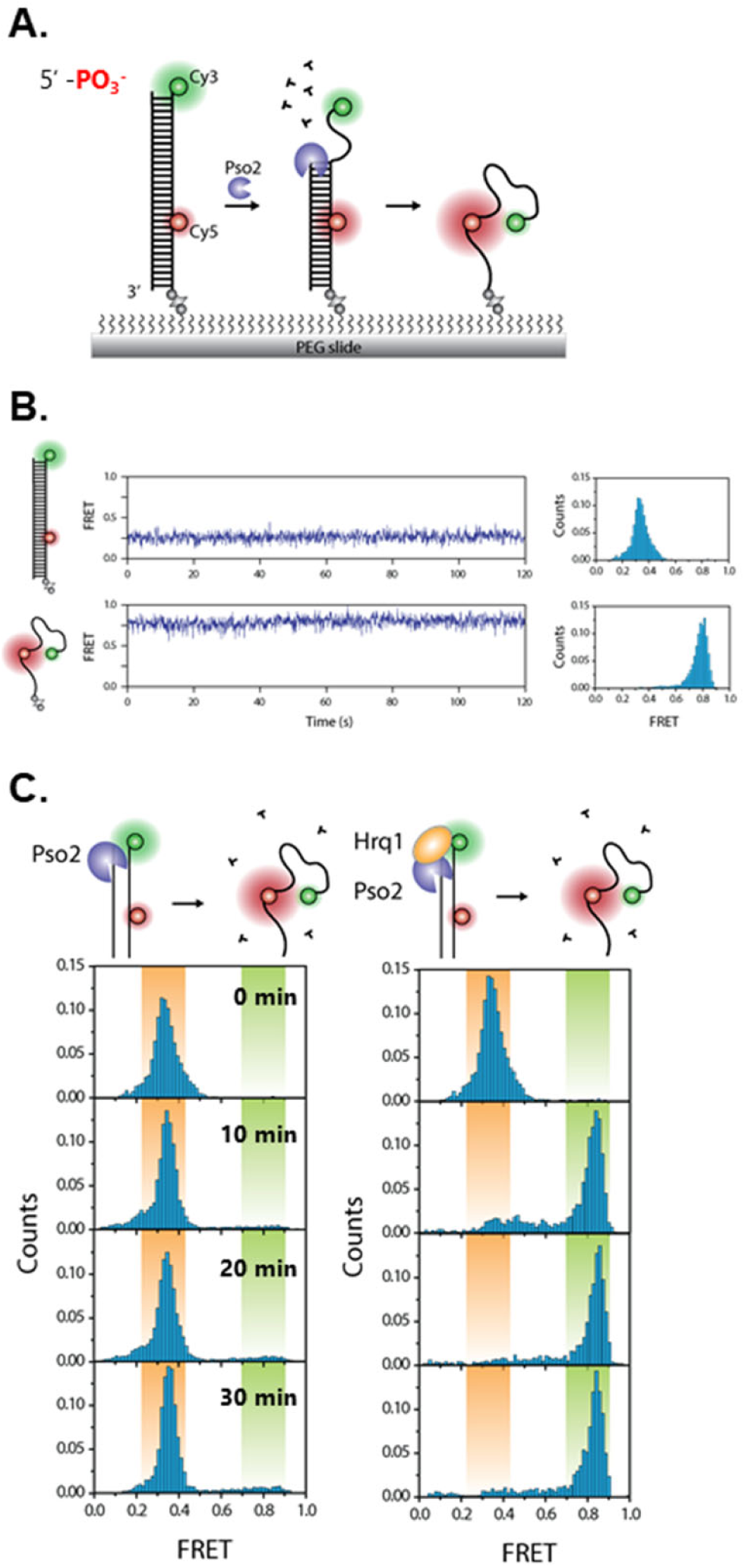
smFRET analysis of the stimulation of Pso2 nuclease activity by Hrq1. **A**. Diagram of the smFRET substrate and the effect of Pso2. Short lengths of dsDNA are rigid, keeping the Cy3 and Cy5 FRET pair distal from one another. Pso2 can digest away the unlabelled DNA strand, yielded flexible ssDNA and allowing the FRET pair to come into proximity. **B**. smFRET signal of the dsDNA substrate and the labelled ssDNA. **C**. Hrq1 stimulates Pso2 nuclease activity. Pso2 alone slowly generates an increase in the FRET signal by degrading the dsDNA substrate, but the addition of Hrq1 greatly increases the speed at which the high FRET signal appears.

To calibrate the smFRET system, we measured FRET from the dsDNA substrate and unannealed ssDNA as undigested and completely digested controls, respectively. The FRET histograms of dsDNA and ssDNA produced sharp peaks at the expected values of 0.32 and 0.80, respectively (Fig. 3B). Next, we applied varying conditions of Pso2 and Hrq1 and collected images in 10-min intervals to measure the FRET change over time. Upon adding 50 nM Pso2 alone, < 10% of molecules shifted to high FRET after 30 min (Fig. 3C) in a Mg^2+^- dependent manner (Fig. S3A). However, longer time courses (> 40 min) of Pso2 nuclease activity revealed that Pso2 was able digest nearly 50% of the substrate (Fig. S4A). When Hrq1 was added without Pso2, the FRET peak remained unchanged in the absence and presence of ATP (Fig. S3A), consistent with the fact that Hrq1 does not bind and unwind blunt dsDNA (14). In contrast, when 50 nM Pso2 and 150 nM Hrq1 were incubated together in our FRET assay, we observed a dramatic increase in high FRET signal within minutes (Fig. 3C). Approximately 76% of molecules were digested in 10 min, with nearly 100% of the DNA digested by 30 min. This stimulation was dependent upon ATP and the catalytic activity of Hrq1, as it was absent when Hrq1-K318A was added to the reaction (data not shown). These smFRET results are highly correlated to our gel-based assays, which also display significantly enhanced Pso2 nuclease activity by Hrq1 (Fig. 2).

Next, we compared the nuclease-helicase coupled activity on undamaged DNA *vs*. a substrate containing a site-specific ICL (XL- DNA). In the absence of Hrq1, Pso2 digested 6% of the undamaged DNA in 10 min, but no activity was detected for Pso2 digestion of XL-DNA (Fig. 4 and S3B), indicating that Pso2 has little to no translesion nuclease activity alone. Even with longer incubation times and in contrast to undamaged DNA, Pso2 was unable to digest a measurable amount of XL-DNA (Fig. S4B). Strikingly, in the presence of Hrq1, 62% of the XL-DNA substrate was digested by Pso2 (Fig. 4A and B).

**Figure 4.**
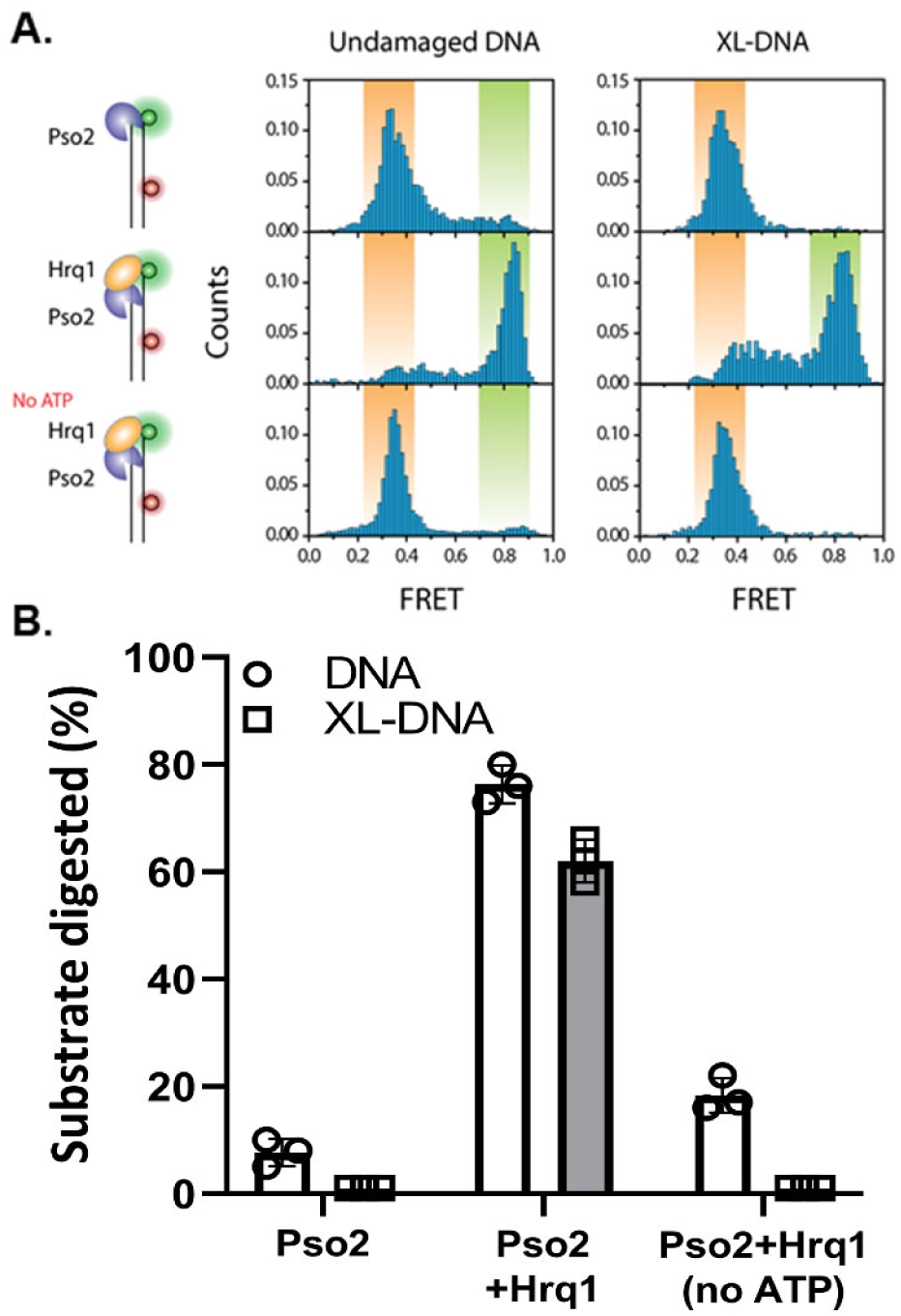
Hrq1 stimulates the translesion nuclease activity of Pso2. **A**. Pso2 lacks translesion nuclease activity in the absence of Hrq1. smFRET analysis of Pso2 activity on undamaged DNA and DNA with a site-specific ICL (XL-DNA) in the absence and presence of Hrq1. ATP is required to observe Pso2 stimulation by Hrq1. **B**. Quantification of the results from A. The graphed bars are the averages of three independent experiments (individual data points shown as open circles), and the error bars are the S.D.

We then asked if the enhanced Pso2 activity requires ATP hydrolysis by Hrq1. To address this question, we conducted the FRET-based nuclease assay in the absence of ATP. Without ATP, Pso2 + Hrq1 digested only 17% of undamaged DNA and no XL-DNA after 10 min (Fig. 4). These values are more comparable to Pso2 in the absence of Hrq1. Similarly, we measured Pso2 nuclease activity in the presence of the ATPase-null Hrq1-K318A mutant, which yielded 8.5% digestion product for undamaged DNA and none for XL-DNA (data not shown), confirming that the ATPase activity of Hrq1 is essential for promoting Pso2 nuclease activity. Taken together, our ensemble biochemical analyses and single molecule data demonstrate that Hrq1 significantly stimulates Pso2 nuclease activity on both undamaged DNA and XL-DNA, and the enhancement requires the catalytic activity of Hrq1.

### Eukaryotic RecQ4 sub-family helicases specifically stimulate Pso2

As stated above, it is not uncommon for helicases and nucleases to function together in DNA repair pathways, as observed with the extensive resection by Sgs1 and Dna2 in HR (35). To determine if the stimulation of Pso2 nuclease activity is a result of general helicase activity or specifically related to Hrq1, we tested RecQ4 sub-family helicases from different species for their ability to stimulate Pso2 nuclease activity. We also tested Sgs1, the other RecQ family helicase in *S. cerevisiae* (homologous to BLM (36)), for its ability to stimulate Pso2 nuclease activity. Interestingly, both Hrq1 and human RECQL4 were able to significantly stimulate Pso2 (Fig. 5A and B). In contrast, the Hrq1 homolog from *Mycobacterium smegmatis*, called SftH (22), was unable to stimulate Pso2, suggesting this phenomenon is specific to eukaryotic RecQ4 helicases. It should be noted that the *M. smegmatis* genome does not encode a homolog of Pso2, so if SftH is involved in ICL repair, it likely functions via a different mechanism than Hrq1/RECQL4.

**Figure 5.**
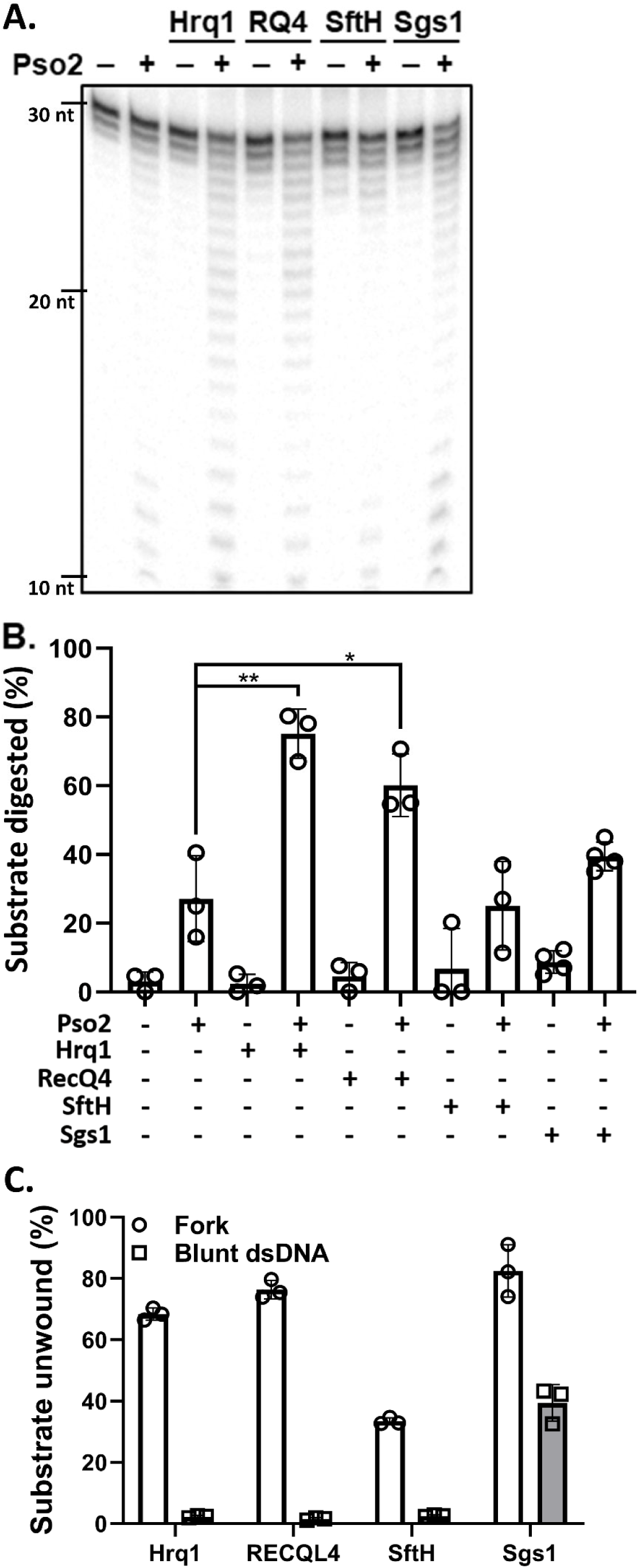
Eukaryotic RecQ4 subfamily helicases specifically stimulate Pso2 nuclease activity. **A**. Pso2 nuclease activity alone and in the presence of recombinant Hrq1, RECQL4, *M. smegmatis* SftH, or Sgs1. The radiolabelled dsDNA substrate was incubated with 50 nM Pso2 and/or 100 nM of the indicated helicase for 30 min, and nuclease products were separated on a denaturing gel and visualized by phosphorimaging. **B**. Quantification of ≥ 3 independent experiments performed as in A. The graphed bars are the averages of the independent experiments (individual data points shown as open circles), and the error bars are the S.D.*, *p* < 0.05 and **, *p* < 0.01. Significant differences were determined by multiple *t*-tests using the Holm-Sidak method, with α = 5% and without assuming a consistent S.D. **C**. Sgs1 unwinds the dsDNA substrate used in nuclease assays. Equimolar amounts of Hrq1, RECQL4, SftH, or Sgs1 were incubated for 30 min with either 2 nM Fork or dsDNA and quantified for helicase activity. The data are plotted as in B.

Sgs1 yielded a level of Pso2 stimulation that was intermediate between SftH and Hrq1/RECQL4 (Fig. 5A and B), though it was not significant (*p* = 0.2). This suggests that the observed synergy with Pso2 is specific to eukaryotic RecQ4 sub-family helicases. However, while *sgs1Δ* cells are also sensitive to ICL damage, they are not epistatic to *hrq1Δ* (25), suggesting that Sgs1 does not function in the Pso2 ICL repair pathway. To investigate this, we performed epistasis analysis with all three of the *hrq1Δ, pso2Δ*, and *sgs1Δ* alleles. As shown in Figures 1 and S1B, we recapitulated the synergistic ICL sensitivity of the *hrq1Δ sgs1Δ* double mutant relative to either of the single mutants (25) and found that the same was true of the *pso2Δsgs1Δ* double and *hrq1Δ pso2 Δsgs1Δ* triple mutants. Thus, *in vivo*, Sgs1 does not participate in the Pso2 ICL repair pathway, even in the absence of Hrq1.

While the results in Figure 5A and B suggest a direct interaction between Pso2 and eukaryotic RecQ4 sub-family helicases, it is also possible that Hrq1 and RECQL4 stimulate Pso2 nuclease activity indirectly by unwinding the DNA probe to provide a more accessible ssDNA substrate for Pso2. While all tested helicases were able to unwind a Y-shaped fork substrate (Fig. 5C), the blunt dsDNA substrate used in the nuclease assays was not unwound by Hrq1, RECQL4, or SftH *in vitro*, while Sgs1 was able to unwind blunt dsDNA. The inability of Hrq1 and RECQL4 to unwind a blunt DNA substrate is consistent with our previous work (14), while the ability of Sgs1 to unwind a similar substrate has been previously reported (37). Thus, the modest amount of Pso2 stimulation by Sgs1 shown in Figure 5A and B could be due to the helicase simply generating ssDNA for Pso2 to degrade, but Hrq1 and RECQL4 must function by a different mechanism. Taken together, these results suggest that Hrq1/RECQL4 and Pso2 directly interact to promote the digestion of ICL-containing lesions.

### Hrq1 and Pso2 likely interact through their N-terminal domains (NTDs)

To determine if Hrq1 and Pso2 directly interact, we performed protein-protein crosslinking using the primary amine-targeting disuccinimidyl sulfoxide (DSSO). Incubation of 500 nM Hrq1 (the ∼130 kDa band) with equimolar amounts of Pso2 in the presence of excess DSSO resulted in a high molecular weight species of approximately 180 kDa in mass as revealed by a western blotting (Fig. 6A). This product could correspond to an Hrq1-Pso2 crosslinked complex. Interestingly, the presence of this 180 kDa product was not dependent on the presence of DNA, but a molar excess of Hrq1 relative to Pso2 increased the amount of this species. This is consistent with our biochemical analyses that suggest a higher Hrq1:Pso2 ratio is optimal for nuclease stimulation (Fig. 2B). However, this association also appeared to be weak, as we were unable to demonstrate it via co-immunoprecipitation of Hrq1 and Pso2 from *S. cerevisiae* lysates, in which both proteins (especially Pso2) exist at very low levels (data not shown). Unfortunately, we also could not artificially elevate the intracellular levels of Pso2 because over-expression led to toxicity (Fig. S5).

**Figure 6.**
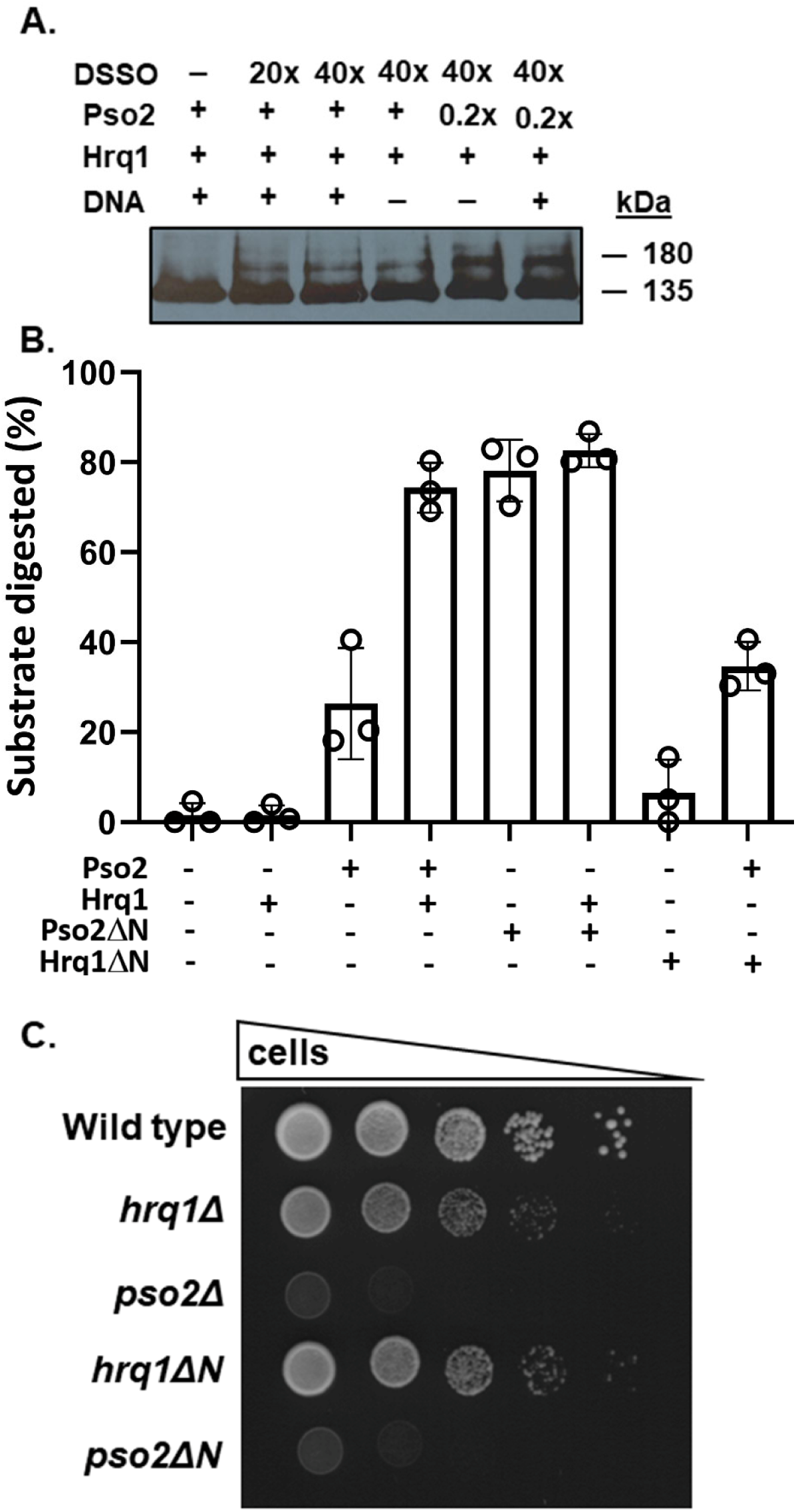
Hrq1 and Pso2 physically interact. **A**. Recombinant Hrq1 and Pso2 can be crosslinked in solution. Molar equivalents of Hrq1 and Pso2 were incubated in the presence or absence of 1 nM poly(dT) 50mer ssDNA and/or the presence or absence of a molar excess (20x or 40x) of the protein-protein crosslinker DSSO. The reactions were then analysed by SDS-PAGE and western blotting with an antibody recognizing the N-terminal 10xHis tag on Hrq1. Hrq1 alone produces a signal at the expected size of ∼135 kDa, but the addition of Pso2 and DSSO yields a band of the approximate size of a Hrq1-Pso2 complex (∼180 kDa). The amount of this slower migrating product increases when Hrq1 is present in a fivefold molar excess of Pso2 (0.2x), consistent with the optimal ratio of Hrq1:Pso2 in nuclease reactions. **B**. Nuclease activity of Pso2 and Pso2ΔN in the absence and presence of Hrq1 or Hrq1ΔN. The Hrq1ΔN truncation mutant fails to stimulate the nuclease activity of full-length Pso2, and Pso2ΔN alone has nuclease activity equivalent to Pso2+Hrq1. The graphed bars are the averages of three independent experiments (individual data points shown as open circles), and the error bars are the S.D.**C**. The *hrq1ΔN* and *pso2ΔN* alleles phenocopy complete deletions of *HRQ1* and *PSO2*. Cells of the indicated genotypes were grown, diluted, and spotted onto YPD + DEB plates as in Figure 1.

We previously hypothesized that the disordered N-terminus of Hrq1 could be an important docking site for protein-protein interactions (14). Similarly, Pso2 is also predicted to possess an unstructured NTD (Fig. 6A). To determine the role of the Pso2 N-terminus in Hrq1-mediated nuclease stimulation, we first compared the nuclease activity of full-length Pso2 to an N-terminal truncation of the first 94 residues (referred to as Pso2ΔN). Interestingly, we found that Pso2ΔN had significantly more nuclease activity than full-length Pso2 (Fig. 6B), suggesting that the Pso2 N-terminus is an autoinhibitory domain. This phenomenon is conserved in the human homolog of Pso2, SNM1A, whose N-terminal truncation is also more active *in vitro* (13). A nuclease-null mutant of Pso2ΔN (Pso2ΔN-H611A) purified identically to Pso2ΔN lacked detectable nuclease activity *in vitro*, indicating that this effect was not due to the presence of a contaminating *E. coli* nuclease (data not shown).

The addition of Hrq1 did not result in stimulation of Pso2ΔN (Fig. 6B), though the highly efficient nuclease activity of the truncation may have already been maximal when used at 20 nM, which is a concentration at which stimulation of full-length Pso2 by Hrq1 can be observed. To test this, we measured the nuclease activity of 2 nM Pso2ΔN, which has a more comparable amount of nuclease activity to 20 nM full-length Pso2 (Fig. S6B). Hrq1 very mildly stimulated Pso2ΔN nuclease activity under these conditions, but not nearly to the levels of full-length Pso2. These data suggest that the Pso2 N-terminus is an autoinhibitory domain that also interacts with Hrq1 to mediate the observed nuclease activity stimulation.

Because *M. smegmatis* SftH was unable to stimulate Pso2 (Fig. 5A and B), we hypothesized that its lack of a large, natively disordered NTD like Hrq1 and RECQL4 may be the reason for this difference. Thus, we measured stimulation of Pso2 by an Hrq1 N- terminal truncation of residues 1-279 (known as Hrq1ΔN). Hrq1ΔN was unable to stimulate Pso2 nuclease activity (Fig. 6B), suggesting the interaction interfaces for both proteins are their disordered N-termini. The *in vivo* activity of these mutants also supports this conclusion as the ICL sensitivity of cells expressing Hrq1ΔN rather than the full-length helicase phenocopied that of a strain completely lacking Hrq1, and *pso2Δ* and *pso2ΔN* cells were also similarly sensitive to ICL damage (Fig. 6C).

## Discussion

Our genetic and biochemical analyses of the Hrq1 and Pso2 interaction suggest that the mechanistic role of Hrq1 in the Pso2 ICL repair pathway is to stimulate Pso2 translesional nuclease activity, facilitating efficient repair of ICLs from multiple sources. Our ensemble and single-molecule biochemistry data indicate that Pso2 alone is unable to digest past an ICL, but Hrq1 facilitates its translesional activity in an ATP-dependent manner. This phenomenon is specific to eukaryotic RecQ4 helicases, likely due to an NTD-to-NTD physical interaction between the helicase and nuclease. Importantly, these results explain why *recql4* mutant cells are sensitive to ICLs, and this DNA repair deficiency may underlie the genomic instability of RECQL4-linked diseases.

### Implications for FA-independent ICL repair across evolution

The FA pathway is the key metazoan ICL repair pathway during S-phase (4), but it has recently been appreciated that FA-independent ICL repair occurs in other contexts, such as during other cell cycle phases and when RNA polymerase encounters an ICL (7,38). In non-metazoans, the FA pathway is also not the dominant ICL repair mechanism or absent altogether. For instance, *S. cerevisiae* predominantly uses the Pso2 repair pathway but contains a rudimentary FA pathway (12,15). However, FA homologs are absent in prokaryotes, and thus, additional solutions to ICL repair exist.

The data presented here and in our previous work (27) indicate that Hrq1 is involved in FA-independent ICL repair. This mirrors what is known about human RECQL4 in ICL repair in that *recql4* mutant cells are sensitive to the ICL-inducing drug cisplatin (26), and RECQL4 does not belong to any of the known FA complementation groups (39,40). Work on the Hrq1 homologs in *Schizosaccharomyces pombe* (41) and *A. thaliana* (24) also supports a role for these helicases in ICL repair, and the more distantly related Hrq1 homolog in *Bacillus subtilis* named MrfA has a role in the repair of MMC-induced lesions (23). Although the function of archaeal RecQ4 helicases (called SftH (22)) is completely unexplored, it is tempting to speculate that they are also involved in ICL repair based on homology to their bacterial and eukaryotic homologs.

The involvement of RecQ4 family helicases in ICL repair across evolution may indicate that the repair pathway in which these helicases function is one of the original mechanisms that developed to combat ICL damage, with the FA pathway evolving much later. ICL repair is a critical genome stability pathway, both in the face of bi-functional exogenous compounds that cause ICLs and endogenous sources of ICL damage. For instance, acetaldehyde is a metabolite produced during ethanolic fermentation that can cause DNA ICLs. It accumulates intracellularly (42), especially when exogenous ethanol concentrations are low (43), and thus ICLs are more apt to occur during the early stage of fermentation. Although we did not test *hrq1Δ* or *pso2Δ* cells for sensitivity to endogenous sources of ICLs, sensitivity of *hrq1* mutants to acetaldehyde-derived ICLs would explain why *HRQ1*/*hrq1Δ* diploids display haploinsufficiency specifically during the early phase of wine fermentation but not later stages when the exogenous ethanol concentration is high (44).

Considering the evolutionary importance of Hrq1-type helicases and ICL repair, it should be noted that not all prokaryotes encode a RecQ4 helicase, with *E. coli* being a prime example. In such organisms, ICL repair is performed by the concerted activities of the UvrABC endonuclease and recombination machinery (45,46). Similarly, not all organisms expressing a RecQ4 sub-family helicase encode a homolog of Pso2/SNM1A, so other proteins that function with RecQ4 helicases in ICL repair in such species await discovery.

### How does Hrq1 stimulate Pso2?

The exact molecular mechanism of Pso2 stimulation by Hrq1 remains unknown. Our analyses of the Pso2 NTD suggest that it is important for ICL repair and likely acts as the interaction interface for Hrq1. It is possible that Hrq1 and Pso2 form a complex in which Hrq1 translocates 3′ → 5′ along the undigested strand, while Pso2 degrades the complementary strand in the 5′ → 3′ direction in a mechanism similar to the Sgs1-Dna2 helicase-nuclease complex used in DSB repair (32). Alternatively, Hrq1 and Pso2 may interact more dynamically when Pso2 stalls. Pso2 alone had very low nuclease activity and processivity (Fig. 2A), though still sufficient for digestion of the short (20-40 nt) excision substrate produced by NER factors in ICL repair. When Pso2 stalls randomly on undamaged DNA or specifically at an ICL, Hrq1 may eject the unproductive nuclease to allow rebinding of Pso2 to continue DNA degradation. A similar model has been proposed for Hrq1 stimulation of telomerase (47).

The requirement of Hrq1 catalytic activity to stimulate Pso2 suggests that Hrq1 does not promote nuclease activity simply by inducing a conformational change in Pso2. However, this could be a component of the mechanism, especially when considering the autoinhibitory nature of the Pso2 NTD. Hrq1 could interact with the Pso2 N-terminus, prohibiting the Pso2 NTD from inhibiting nuclease activity. While our data indicate that this is not the sole explanation for Hrq1-mediated stimulation of Pso2, we cannot exclude this proposed phenomenon as a component of the mechanism.

Pso2 has an endonucleolytic activity under certain *in vitro* conditions (18), so Hrq1 may appear to stimulate Pso2 DNA digestion simply by unwinding dsDNA into ssDNA to facilitate endonucleoytic cleavage of a single-stranded substrate. Our data showing that Hrq1 and RECQL4 cannot unwind blunt dsDNA (Fig. 5C) but both stimulate Pso2 nuclease activity on the same substrate (Fig. 5A and B) argue against this. According to the model in the field ((15) and Fig. 7), Pso2 acts after the NER machinery makes incisions on one strand of DNA on either side of the ICL. *In vitro*, Pso2 can load at a nick to degrade DNA in the 5′ → 3′ direction (18). Therefore, Pso2 likely does not require endonuclease activity during ICL repair, when it presumably loads at the 5′ nick in the DNA upstream of the ICL. However, we have not tested the ability of Hrq1 to stimulate Pso2 endonuclease activity nor if Hrq1 can load at a nick to initiate DNA unwinding. Such experiments are ongoing because specifically identifying how Hrq1 stimulates Pso2 nuclease activity will be important when translating this model to human RECQL4 and SNM1A.

**Figure 7.**
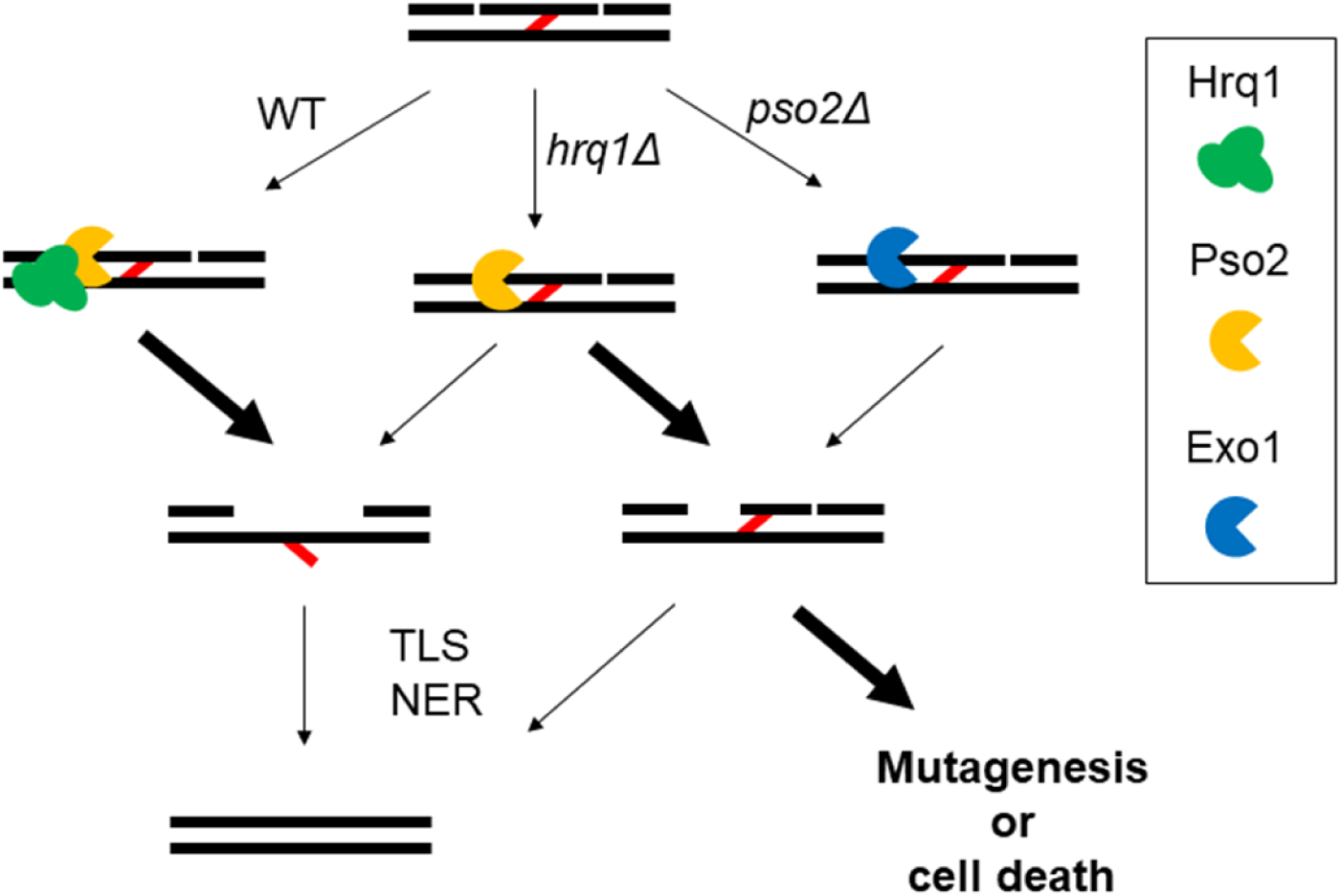
Model of ICL repair in WT, *hrq1Δ*, and *pso2Δ* cells. In the first step of ICL repair, the NER machinery cuts one strand of DNA on either side of the ICL. Then, in WT cells, Hrq1 and Pso2 are recruited to the lesion to digest away the incised strand, leaving an adducted base. Translesion synthesis (TLS) polymerases fill the gap, and NER removes the adducted base. In *hrq1Δ* cells, Pso2 is still recruited to the ICL, but its poor translesion nuclease activity in the absence of Hrq1 yields some amount of incompletely processed substrates, which can lead to mutagenesis or cell death. In cells lacking Pso2, other nucleases (*e*.*g*., Exo1) may be recruited to ICLs, but there less optimal activity on such substrates can also lead to mutagenesis or cell death.

### Regulation of Pso2 nuclease activity

Why does Hrq1 need to stimulate Pso2 for appropriate ICL resistance? Pso2 expression is extremely low, and ICL damage only results in a modest induction of Pso2 (48). The overexpression of Pso2 in yeast is extremely toxic (Fig. S5), and the Pso2 N-terminus autoinhibits nuclease activity (Fig. 6B), providing another regulatory mechanism to prevent rampant Pso2 digestion. Thus, Pso2 expression is in a delicate balance between being sufficient for ICL repair and nucleotoxic. Coupling Hrq1 to Pso2 in ICL repair may allow for modulated, site-specific nuclease activity. In this scenario, Hrq1 and Pso2 are recruited to ICLs via an unknown mechanism, and low levels of Pso2 are sufficient for ICL degradation, as Hrq1 is present to stimulate translesion exonuclease activity (Fig. 7). This scheme allows for cells to maintain low Pso2 levels but still have the appropriate amount of nuclease activity at the lesion. In the absence of Hrq1, Pso2 is still recruited to ICLs and can aid in their repair, but the process is less efficient, accounting for the mild ICL sensitivity of *hrq1Δ* cells (Fig. 1 and S1). Finally, in the absence of Pso2 itself, other pathways can repair some amount of the lesions (*e*.*g*., the proto-FA pathway (15)), but many ICLs likely persist, resulting in mutagenesis and death (Fig. 7).

These data provide evidence of a novel role for RecQ4 helicases in ICL repair and further support the model that translesion nuclease activity by Pso2 is an important step in ICL repair. Because components of this mechanism are likely conserved in all domains of life, future work will be required to identify new proteins that facilitate RecQ4-medicated ICL repair and determine how RecQ4 helicases operate across evolution. How cells utilize such a diverse set of tools for ICL repair is likely determined by the molecular context in which the lesions are encountered, and pathway choice is an important but largely unexplored element of ICL repair.

## Experimental procedures

### Reagents, nucleotides, and oligonucleotides

γ[^32^P]-ATP and α[^32^P]-dCTP were purchased from PerkinElmer (Waltham, MA). Unlabelled ATP was purchased from DOT Scientific (Burton, MI), and all oligonucleotides were synthesized by IDT (Coralville, IA) and are listed in Supplemental Table 1.

### DNA inter-strand crosslinker sensitivity analysis

The sensitivity of *S. cerevisiae* mutants (Table S2) to DNA damaging agents was analysed as described (14). Briefly, cells were grown overnight in 1% yeast extract, 2% peptone, and 2% dextrose (YPD) medium with aeration and diluted to an optical density at 660 nm (OD_660_) of 1 in sterile water. Cells were then serially diluted 10-fold to 10^−4^, and 5 µL of each dilution was spotted onto YPD plates lacking or containing the indicated drug. For MMC and DEB, plates contained 50 μg/mL of the respective drug, unless otherwise indicated.

Plates containing 20 µg/mL 8-MOP were treated with 365 nm UVA (Sylvania fluorescent lamp) in a dark box for 30 min after cells were spotted to activate the ICL reaction. MMS was used at 0.03%. All plates were incubated in the dark for 2 days at 30°C and imaged on a flat-bed scanner. The strains were constructed in the wild type YPH499 background (*MATa, ura3- 52, lys2-801_amber, ade2-101_ochre, trp1Δ63, his3Δ200, leu2Δ1*) (49) using standard methods.

### Protein expression and purification

The *S. cerevisiae* Pso2 expression vector harbouring a C-terminal 6xHis-tag was kindly provided by Murray Junop (Schulich School of Medicine and Dentistry, Western University) (18). Pso2-6xHis was expressed in Rosetta 2(DE3) pLysS (Novagen) cells by growing cultures to an OD_600_ of 0.6 at 37°C followed by induction with 1 mM isopropyl β-D-1- thiogalactopyranoside for 4 h at 30°C. Cells were harvested by centrifugation at 5000 rpm for 10 min at 4°C, and the cell pellet was frozen at −80°C. The frozen cell mass was thawed in Resuspension Buffer (50 mM NaHEPES, pH 7, 50 mM NaCl, 5% glycerol, and 2 mM DTT) supplemented with fresh protease inhibitor mix and 20 μg/mL DNase I. Lysis was performed using several passes through a cell cracker, and the lysate was clarified by centrifugation at 14,000 rpm for 30 min at 4°C. Clarified lysate was loaded onto a gravity column containing 1 mL HIS-Select Nickel Affinity Gel (Sigma) pre-equilibrated with Resuspension Buffer. The column was washed with 5 column volumes (CVs) of Resuspension Buffer and 5 CVs of Resuspension Buffer supplemented with 5 mM ATP. Pso2 was eluted with Resuspension Buffer supplemented with 100 mM imidazole, and the Pso2-containing fractions were pooled. The eluate was then loaded onto 1 mL HiTrap Heparin HP column (GE Healthcare) and washed with 10 CVs Heparin Buffer (50 mM NaHEPES, pH 7, 50 mM NaCl, 5% glycerol, and 2 mM DTT). Pso2 was eluted via a 20-CV linear salt gradient with Heparin Buffer from 50 mM to 1 M NaCl. Pooled Pso2 fractions were concentrated and buffer exchanged into storage buffer (25 mM NaHEPES, pH 7.6, 30% glycerol, 300 mM NaOAc pH 7.6, 25 mM NaCl, 5 mM MgOAc, 1 mM DTT, and 0.01% Tween- 20). Pso2ΔN and the catalytically inactive Pso2-H611A and Pso2ΔN-H611A mutants were purified identically to wild-type.

Expression and purification of Hrq1 and RECQL4 were described previously (14). Sgs1 was a generous gift from Hengyao Niu (Indiana University) and Petr Cejka (Università della Svizzera italiana).

Hrq1ΔN was purified similarly to the wild-type Hrq1 used in (25). Briefly, a plasmid harboring the 3xStrep-Hrq1ΔN-6xHis construct was transformed into Rosetta 2(DE3) pLysS cells and expressed overnight in autoinduction media (50). Cells were lysed in Hrq1ΔN Resuspension Buffer (50 mM NaHEPES,pH 8, 5% glycerol, 150 mM NaOAc, pH8, 5 mM MgOAc, and 0.05% Tween-20) as in the Pso2 purification, and clarified lysate was loaded onto a 1 mL StrepTrap column. The column was washed with 20 CVs Hrq1ΔN Resuspension Buffer supplemented with 600 mM NaOAc, followed by a wash with Hrq1ΔN Resuspension Buffer supplemented with 5 mM ATP. Protein was eluted in buffer containing 2.5 mM desthiobiotin. Protein-containing fractions were pooled and loaded onto a 0.5-mL His60 column. The column was washed with 10 CVs Hrq1ΔN Resuspension Buffer containing 10 mM imidazole, followed by a 1-CV wash each with buffer supplemented with 50 and 100 mM imidazole. Protein was eluted in Hrq1ΔN Resuspension Buffer supplemented with 500 mM imidazole and stored as for Pso2. *Mycobacterium smegmatis* SftH was purified identically.

Further details concerning the construction of expression vectors and protein purification are available upon request. Mass spectrometry analysis of all protein preparations demonstrated that they were not contaminated by *E. coli* helicases or nucleases. Representative stained SDS-PAGE gel images of the recombinant proteins are shown in Figure S7.

### Preparation of DNA substrates

The uncrosslinked double-stranded (ds)DNA substrate was prepared by annealing equimolar amounts of MB1614 to MB1461 in Annealing Buffer (20 mM Tris-HCl, pH 8, 4% glycerol, 0.1 mM EDTA, 40 µg/mL BSA, 10 mM DTT, and 10 mM MgOAc) overnight at 37°C (51) (Table S1). Because Pso2 nuclease activity is greatly stimulated by a 5′-phosphate, only the digested strand (MB1614) was phosphorylated by IDT. The substrate was designed such that the digested strand is 6 nt shorter than the undigested strand to allow 3′ fill-in by Klenow Fragment (3′-5′ exo-; NEB) with α[^32^P]-dCTP and cold dATP, dTTP, and dGTP for 30 min at 37°C. After labelling, cold dCTP was also added for another 30 min at 37°C to facilitate complete fill-in to yield a 30-bp blunt dsDNA. The nuclease substrate was separated from unincorporated dNTPs using an illustra ProbeQuant G-50 micro column (GE Healthcare) according to the manufacturer’s instructions.

The ICL-containing substrate was prepared using spontaneous crosslink formation from an abasic site as previously reported (33). Oligonucleotides MB1599 and MB1600 were annealed by heating to 95°C for 5 min followed by slow cooling to room temperature overnight. The digested strand (MB1599) was 5′ phosphorylated as above and contained a deoxyuracil (dU) 7-nt from the 5′ end. The substrate was then treated with 50 U of uracil DNA glycosylase (NEB) for 2 h at 37°C to form the abasic site. The DNA was phenol/chloroform extracted and precipitated with 10% 3 M NaOAc, pH 5.2, and five volumes of 100% ethanol. Precipitated DNA was stored at −20°C for 1 h and pelleted at 15,000 rpm for 30 min at 4°C. The supernatant was removed, and the pellet was washed twice with cold 80% ethanol. After the last wash was completely removed, the DNA was resuspended in 50 mM NaHEPES, pH 7, and 100 mM NaCl and stored in the dark for 5 days with gentle agitation to form the ICL. Once the crosslink was formed, the substrate was labelled with α[^32^P]-dCTP as above. The DNA was heated at 95°C for 10 min to denature any uncrosslinked substrate, and the entire sample was loaded onto a 20% 19:1 acrylamide:bis-acrylamide 6 M urea denaturing gel and run in 1x TBE buffer (90 mM Tris-HCl pH 8.0, 90 mM boric acid, and 2 mM EDTA, pH 8.0) at 10 V/cm. The gel was exposed to classic blue autoradiography film to identify the slower migrating ICL-containing dsDNA, which was gel extracted into 0.5x TBE buffer overnight and precipitated as above. The crosslinked substrate was finally resuspended in H_2_O.

To make forked DNA for helicase assays, the top strand (MB733) was 5′ labelled with ϒ[^32^P]- ATP and T4 polynucleotide kinase (T4 PNK; NEB) for 1 h at 37°C and cleaned up with a G-50 micro column as above. Equimolar cold bottom strand (MB734) was incubated with labelled top strand overnight at 37°C in Annealing Buffer. Helicase assays using blunt dsDNA were performed with the undamaged nuclease assay substrate described above (MB1614 annealed to MB1461).

### Gel-based nuclease assays

Nuclease assays were performed for 30 min at 30°C with the indicated protein concentration in Nuclease Buffer (20 mM Tris-acetate, pH 7.6, 50 mM NaOAc, pH 7.6, 7.5 mM MgOAc, and 0.01% Tween-20) with 1 nM labelled DNA substrate. For nuclease assays involving helicases, 5 mM ATP was also included unless stated otherwise. Reactions were stopped with the addition of Loading Dye (95% formamide and 0.02% bromophenol blue) and by heating at 95°C for 5 min. Reactions were loaded onto 20% 19:1 acrylamide:bis-acrylamide 6 M urea denaturing gels in 1x TBE buffer and run at 2400 V for 90 min. Gels were dried under vacuum, imaged using a Typhoon FLA 9500, and quantified using ImageQuant 5.2.

### smFRET nuclease assays

Single molecule assays were performed by using a home-built prism type total internal reflection fluorescence (TIRF) microscope at room temperature (24 ± 1°C). The microscope setup, slide preparation, and DNA immobilization were performed as described (34). The experimental protocol and flow setup were slightly modified from a previously published method (52). The MB1621 DNA oligonucleotide was purchased from IDT with 5’-Cy3, 3’-biotin, and an internal amine modification (Supplementary Table 1). The amine modification was used for Cy5 labeling as described (53). The labelled oligonucleotide was annealed to MB1620 or MB1622 to make the ICL-containing or undamaged substrate, respectively. The abasic site crosslink was formed as above.

## Acknowledgements

The buffer used for Pso2 nuclease activity measurement was prepared freshly by mixing an oxygen scavenging system (1 mg/mL glucose oxidase, 0.8% v/v glucose, ∼10 mM Trolox, and 0.03 mg/mL catalase) before taking each single molecule image. A solid-state 532 nm laser was used for FRET measurement. Images were recorded with a time resolution of 100 ms and analysed by Matlab. FRET efficiency (E) was calculated by the equation:

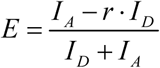

I_D_ and I_A_ are the intensities of Cy3 (donor) and Cy5 (acceptor). *r* is the correction factor for donor leakage, which is 0.15 for our system. Each FRET histogram was generated by collecting FRET values from at least 6000 molecules taken over 15∼20 movies. The FRET histograms were fit with a Gaussian distribution function. For nuclease activity, 50 nM Pso2 and 150 nM Hrq1 (or Hrq1-K318A) were used in all of the experiments, and each reaction was terminated by 0.1% SDS treatment and flushed with TE (90 mM Tris-HCl, pH 8, and 2 mM EDTA, pH 8) to remove protein.

## Protein-protein crosslinking

Disuccinimidyl sulfoxide (DSSO) was resuspended in DMSO to the desired concentration. Reactions contained 500 nM Hrq1, 500 or 100 nM Pso2, and a molar excess of DSSO as indicated. After incubation for 30 min at room temperature, reactions were quenched with 0.5 µL 1 M Tris-HCl, pH 8, and run on 10% SDS-PAGE gels. The proteins were transferred to nitrocellulose and probed with an α-His primary antibody and HRP-conjugated goat α-mouse secondary antibody using standard methods.

## Data availability

Data not shown in this article are available upon request from Matthew L. Bochman.

We thank Drs. Hengyao Niu and Petr Cejka for the gift of recombinant Sgs1, Dr. Murray Junop for providing the Pso2 expression plasmid, Ms. Jade Katinas for purifying SftH, and members of the Bochman lab for useful discussions and critically reading this manuscript.

## Funding and additional information

This work was supported by funds from the College of Arts and Sciences, Indiana University (to MLB); the Indiana University Collaborative Research Grant fund of the Office of the Vice President for Research (to MLB and YT); The American Cancer Society (RSG-16-180-01-DMC to MLB); the National Institutes of Health (1R35GM133437 to MLB, GM115631 to SM, and GM111695 to YT); and the National Science Foundation (MCB-1157688 to YT). Funding for open access charge: National Institutes of Health 1R35GM133437. The content is solely the responsibility of the authors and does not necessarily represent the official views of the National Institutes of Health.

## Conflict of Interest

The authors declare that they have no conflicts of interest.

## Abbreviations

ICL: inter-strand crosslink
FA: Fanconi anemia
DSB: double-strand break
HR: homologous recombination
NER: nucleotide excision repair
ssDNA: single-stranded DNA
MMC: mitomycin C
8-MOP: 8-methoxypsoralen
DEB: diepoxybutane
MMS: methylmethansulfonate
smFRET: single-molecule Förster resonance energy transfer
XL-DNA: ICL-containing DNA
DSSO: disuccinimidyl sulfoxide
NTD: N-terminal domain
dsDNA: double-stranded DNA
TIRF: total internal reflection fluorescence
OD_660_: optical density at 660 nm
S.D.: standard deviation
TLS: translesion synthesis

## Notes

### Competing Interest Statement

The authors have declared no competing interest.

### Summary of Updates

Updated text and figures

