## Supporting information for "The Hrq1 helicase stimulates Pso2 translesion nuclease activity to promote DNA inter-strand crosslink repair"

### Supporting experimental procedures

### Pso2 overexpression in *Saccharomyces cerevisiae*

Pso2 was cloned under the control of the galactose-inducible promoter in pESC-URA and transformed into cells deleted for *pso2*. Empty vector strains were also constructed. Overnight cultures were grown in uracil dropout medium supplemented with 2% raffinose. Cells were pelleted and washed with sterile $H_2O$ before being added to a 96-well plate at an $OD_{660}$ of 0.01 in uracil dropout medium supplemented with either 2% glucose or galactose. Cells were incubated at 30°C with vigorous shaking in a plate reader for 48 h, and $OD_{660}$ readings were taken every 15 min. The resulting growth curves were used to calculate the mean $OD_{660}$ for each condition. The mean values were taken from ≥ 3 independent experiments and averaged. The reported mean $OD_{660}$ values were determined by dividing Pso2 overexpression strains by the mean $OD_{660}$ of the empty vector strains in either glucose (repressed expression) or galactose (overexpression) as indicated.

### Helicase assays

Helicase assays were performed as reported (14). Briefly, 2 nM labelled DNA substrate was incubated with the indicated helicase concentration in Nuclease Buffer with 5 mM ATP for 30 min at 30°C. For RECQL4 helicase assays, 15 nM cold ssDNA trap was added to capture the unwound product. The trap was added last along with ATP to start the reaction. Assays were stopped with 1x Stop-Load dye (5% glycerol, 20 mM EDTA, 0.05% SDS, and 0.25% bromophenol blue) and loaded onto 12% 19:1 acrylamide:bis-acrylamide nondenaturing gels and run in 1x TBE at 10 V/cm. Gels were dried under vacuum, imaged using a Typhoon FLA 9500, and quantified using ImageQuant 5.2.

**Supporting figures**

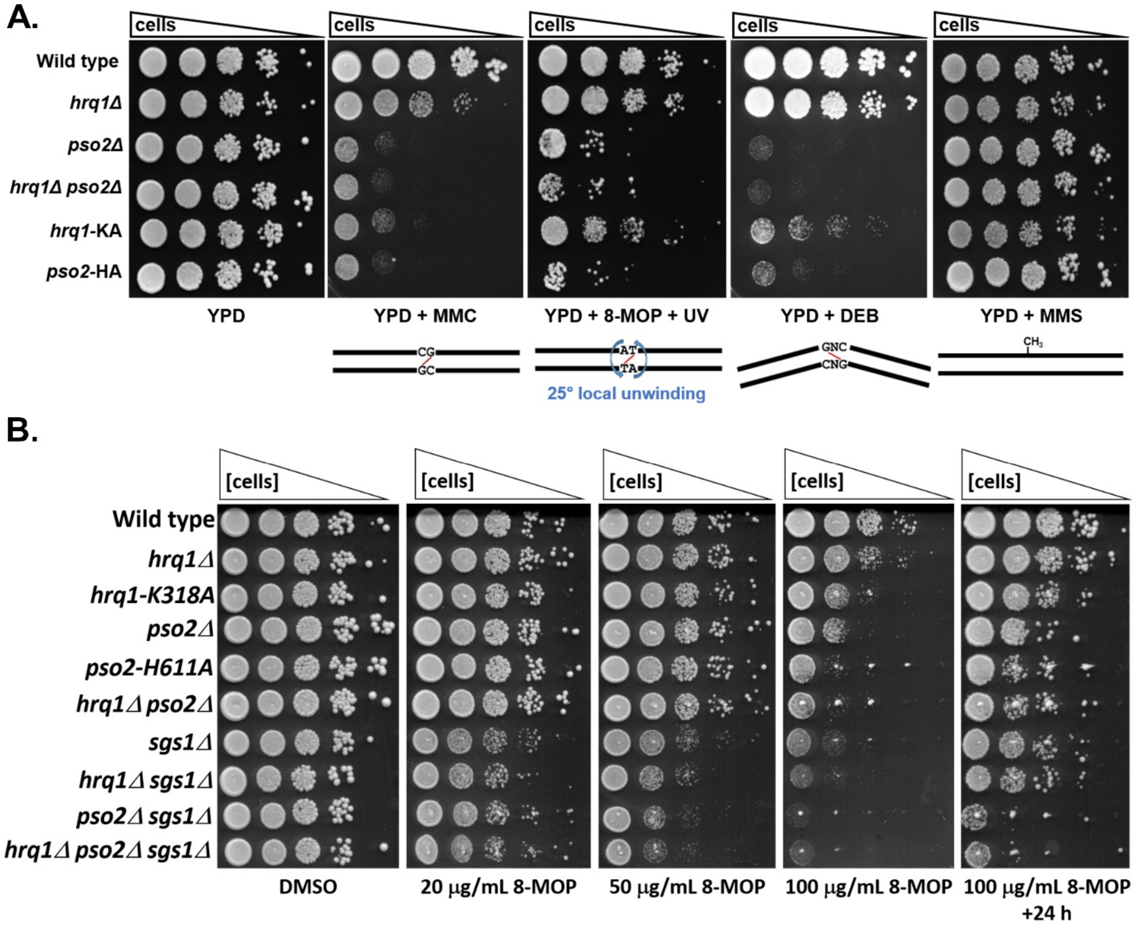


**Figure S1.** Mutation of *HRQ1* and *PSO2* sensitizes cells to ICL damage. **A.** Saturated overnight cultures of strains with the genotypes indicated on the left were diluted to $OD_{660} = 1.0$ and then further serially diluted 10-fold to $10^{-4}$. Equal volumes of each dilution were then spotted onto rich medium (YPD) or rich medium supplemented with MMC, 8-MOP, DEB, or MMS. The plates containing 8-MOP were also exposed to UVA to activate the 8-MOP for crosslinking. The effect of the DNA damaging agent is diagrammed below each plate: MMC does not deform the DNA backbone, 8-MOP + UVA results in ~25° of local unwinding of the DNA around the lesion, DEB kinks the DNA backbone, and MMS alkylates the DNA. **B.** 8-MOP + UVA sensitivity of *hrq1*, *pso2*, and *sgs1* mutants. The assay was performed as described in Figure 1A, and the DMSO control plate from Figure 1 is shown again for ease of comparison. These results are representative of ≥ 3 independent experiments.

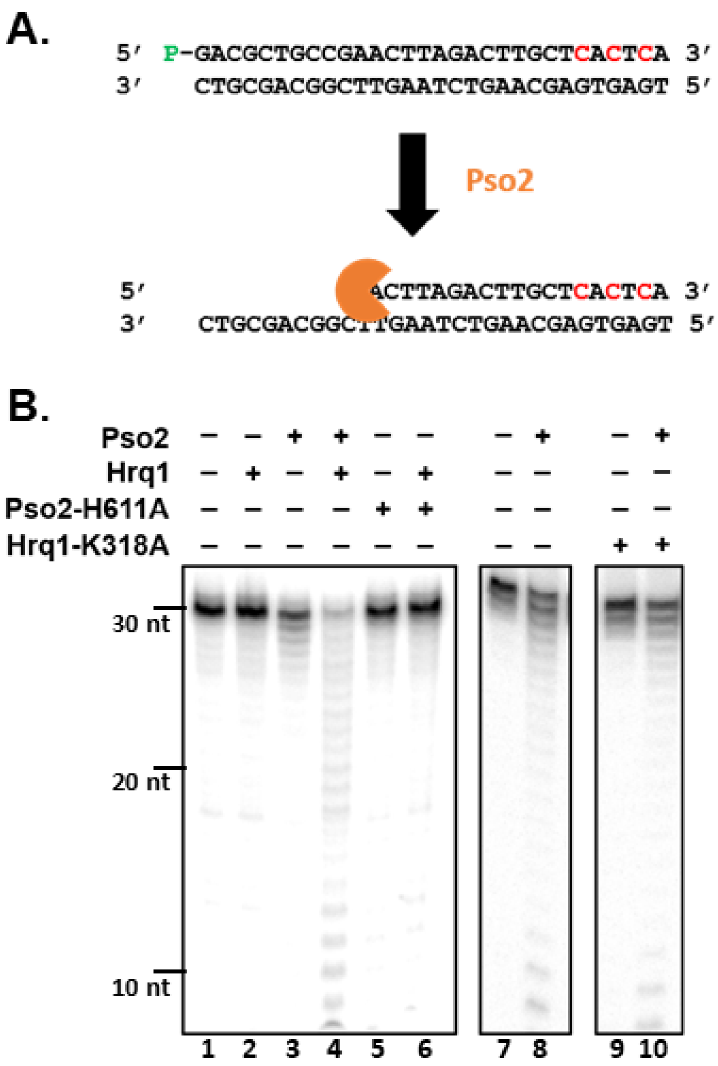


**Figure S2.** Representative gel images of Hrq1-mediated Pso2 stimulation. **A.** Schematic of gel-based nuclease assay. The digested strand is 5′ phosphorylated and labelled on the 3′ end via Klenow fill-in (red nucleotides). Pso2 digestion is measured by the smaller bands as observed on denaturing gels. **B.** Nuclease activity of Pso2 or nuclease-inactive Pso2-H611A in the presence or absence of Hrq1 or helicase-inactive Hrq1-K318A. The indicated combination of enzymes (50 nM nuclease and 150 nM helicase) were incubated with dsDNA for 30 min. Lanes 1-6 were run on the same gel, whereas lanes 7-10 were identical conditions run on a separate gel (with intervening lanes removed for simplicity). Quantification of similar data from at least three independent experiments is reported in Figure 2C.

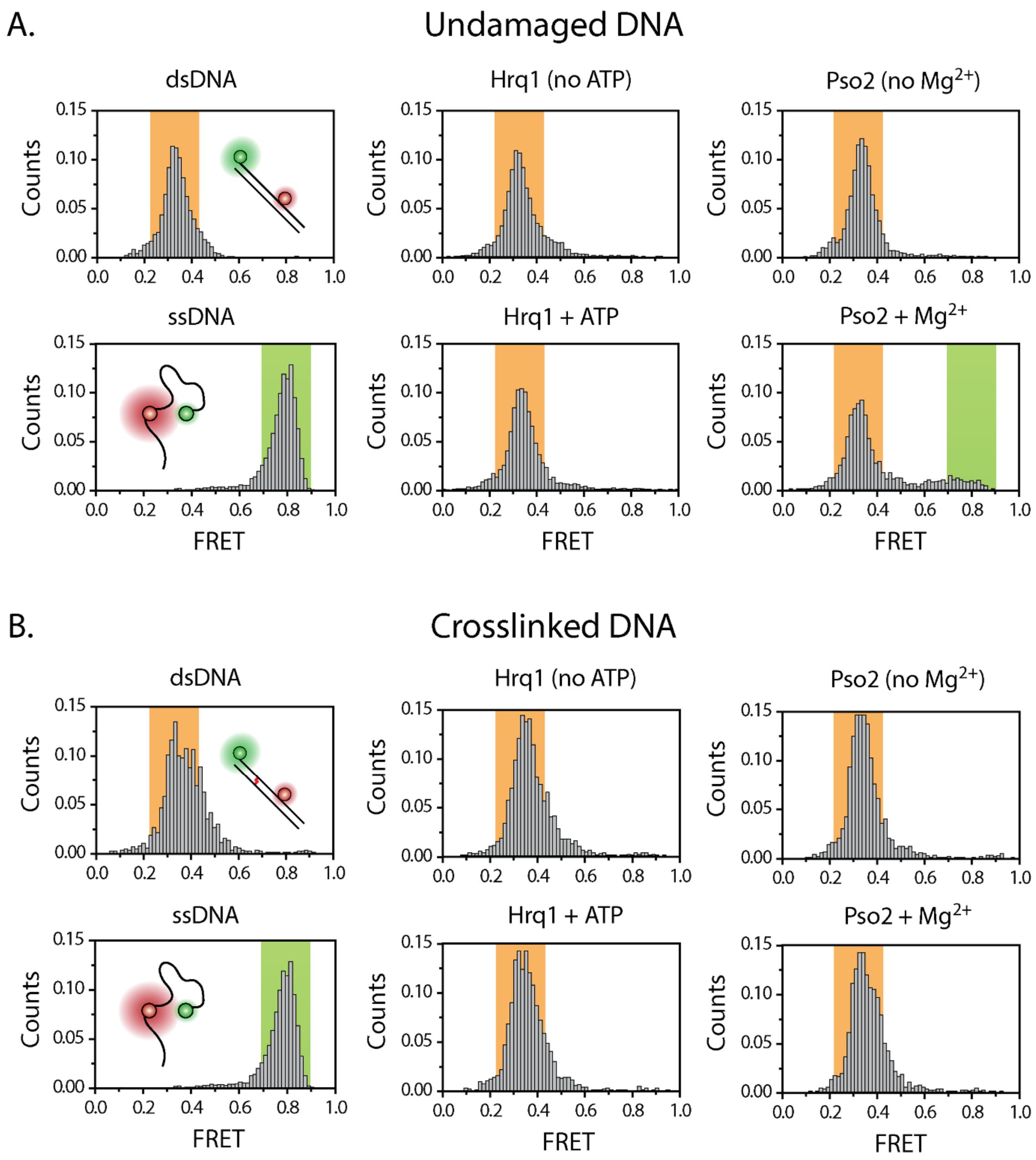


**Figure S3.** Control smFRET nuclease assays. **A.** smFRET analysis on dsDNA under various conditions. The data used to plot the ssDNA and dsDNA histograms were collected in the absence of protein to determine the FRET values of digested and undigested substrate, respectively. Hrq1 +/- ATP had no effect on the substrate, as expected by previous experiments demonstrating a lack of Hrq1 helicase activity on similar blunt dsDNA substrates (14). Pso2 requires the presence of $Mg^{2+}$ for nuclease activity. **B.** The same control experiments from A) were performed with XL-DNA.

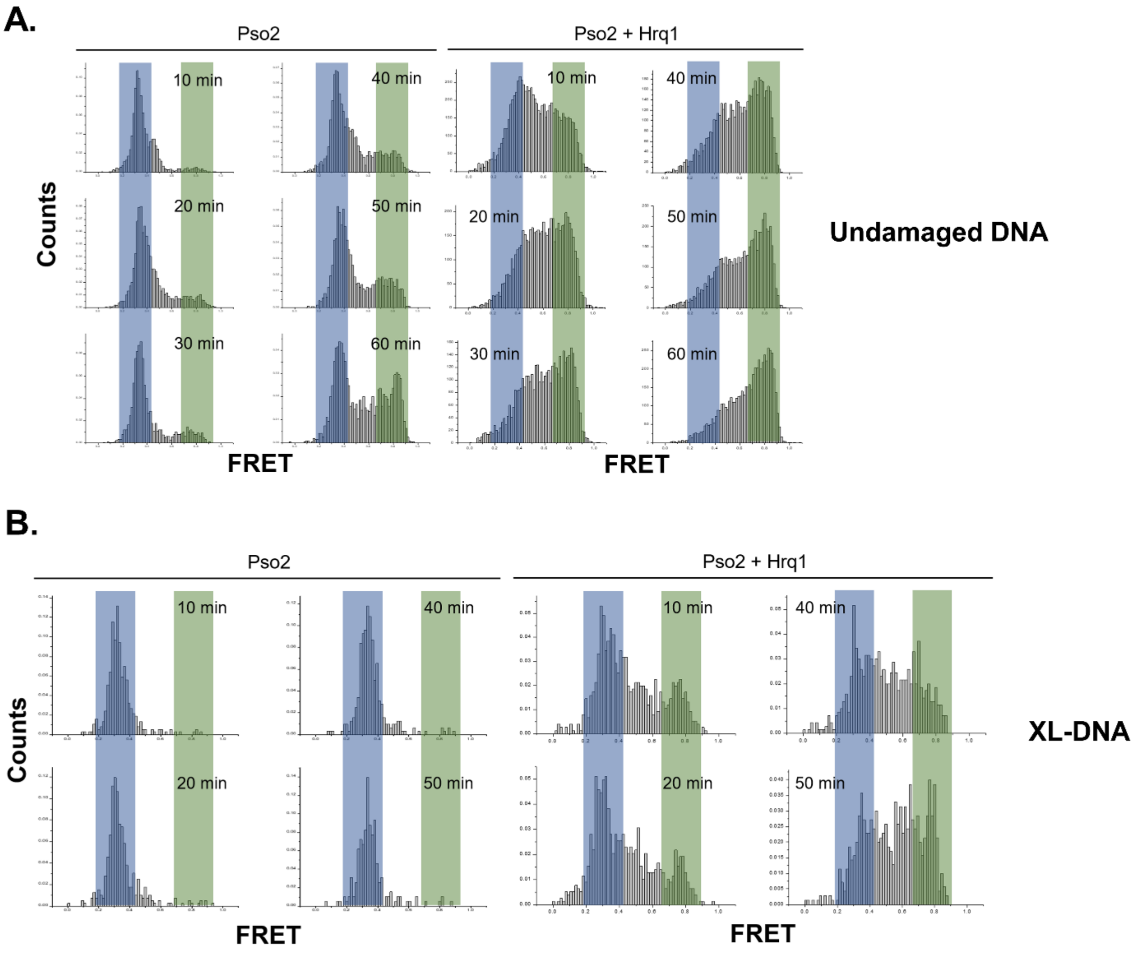


**Figure S4. A.** smFRET analysis of Pso2 nuclease activity on undamaged dsDNA without and with Hrq1. **B.** smFRET analysis of Pso2 nuclease activity on XL-DNA without and with Hrq1.

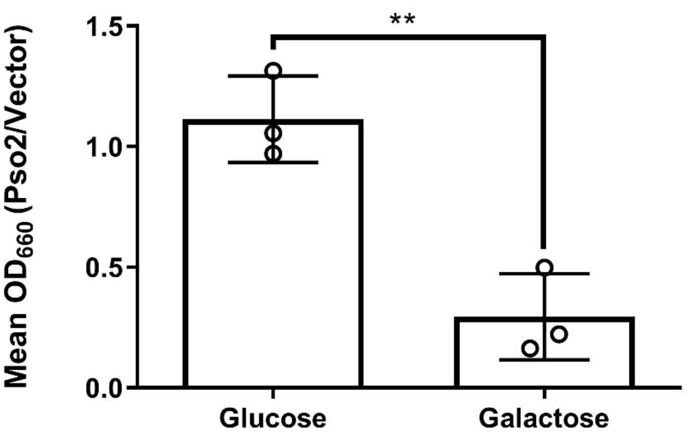


**Figure S5.** Pso2 overexpression is toxic. Growth of cells harbouring a Pso2 overexpression vector in glucose, where Pso2 expression is repressed, is similar to that of the empty vector (Mean $OD_{660}$ (Pso2/vector) of 1). Overexpression of Pso2 via galactose induction severely inhibits cell growth. The graphed bars are the averages of three independent experiments (individual data points shown as open circles), and the error bars are the S.D. ** $p < 0.001$.

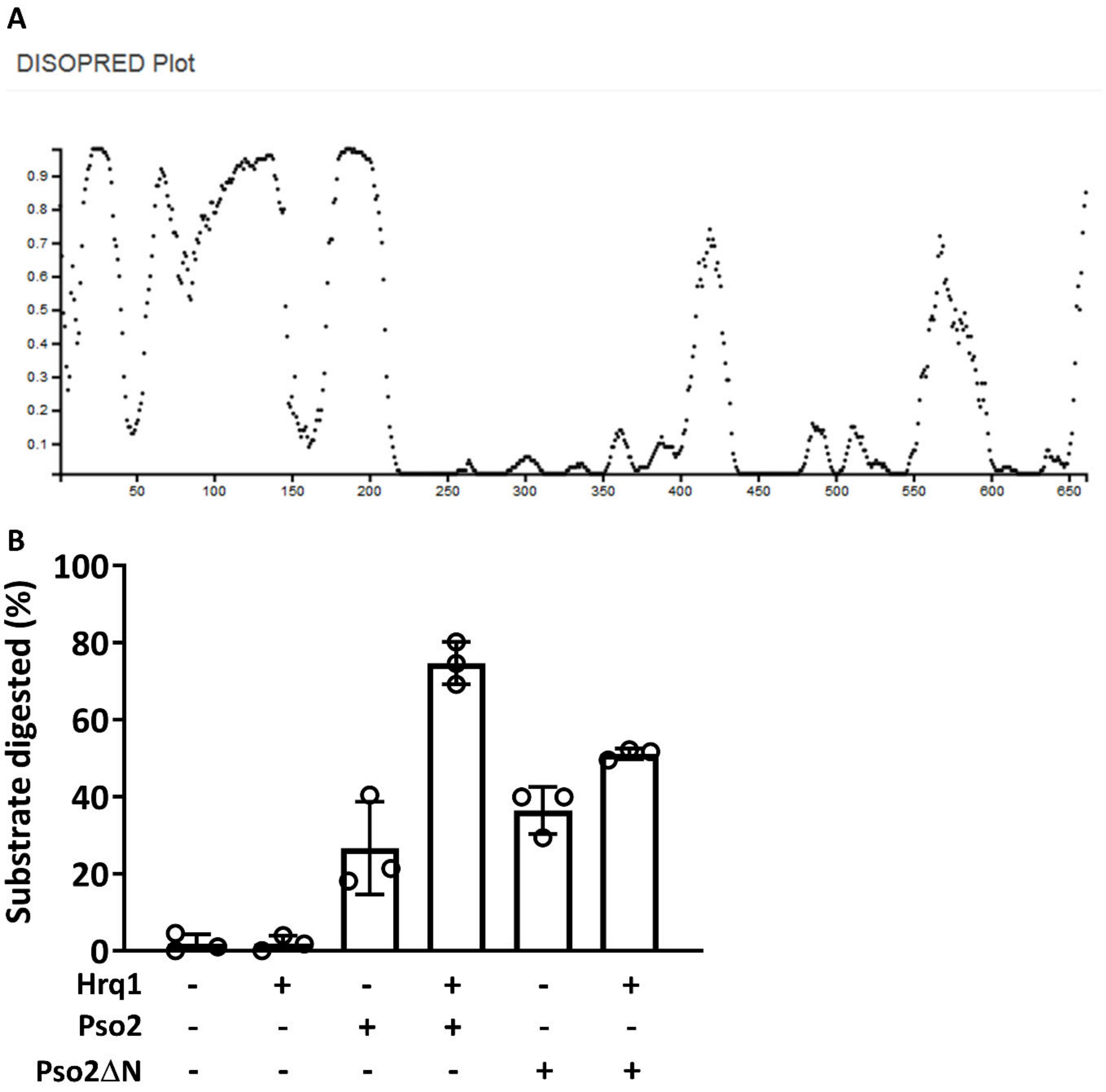


**Figure S6.** The Pso2 N-terminus is an autoinhibitory domain. **A.** A DISOPRED profile for Pso2, as analysed by the UCL Department of Computer Science web server (http://bioinf.cs.ucl.ac.uk/psipred/?disopred=1) indicates that the N-terminal domain of Pso2 (aa 1~210) is predicted to be natively disordered. **B.** Hrq1 is unable to maximally stimulate Pso2ΔN. Pso2ΔN (50 nM) nearly completely digested the dsDNA substrate in Figure 6B, but here, 2 nM Pso2ΔN has comparable nuclease activity to 50 nM full-length Pso2. The addition of Hrq1 yielded very mild stimulation of Pso2ΔN nuclease activity at this concentration, suggesting that the N-terminus of Pso2 is required for Hrq1-mediated stimulation. The graphed bars are the averages of three independent experiments (individual data points shown as open circles), and the error bars are the S.D.

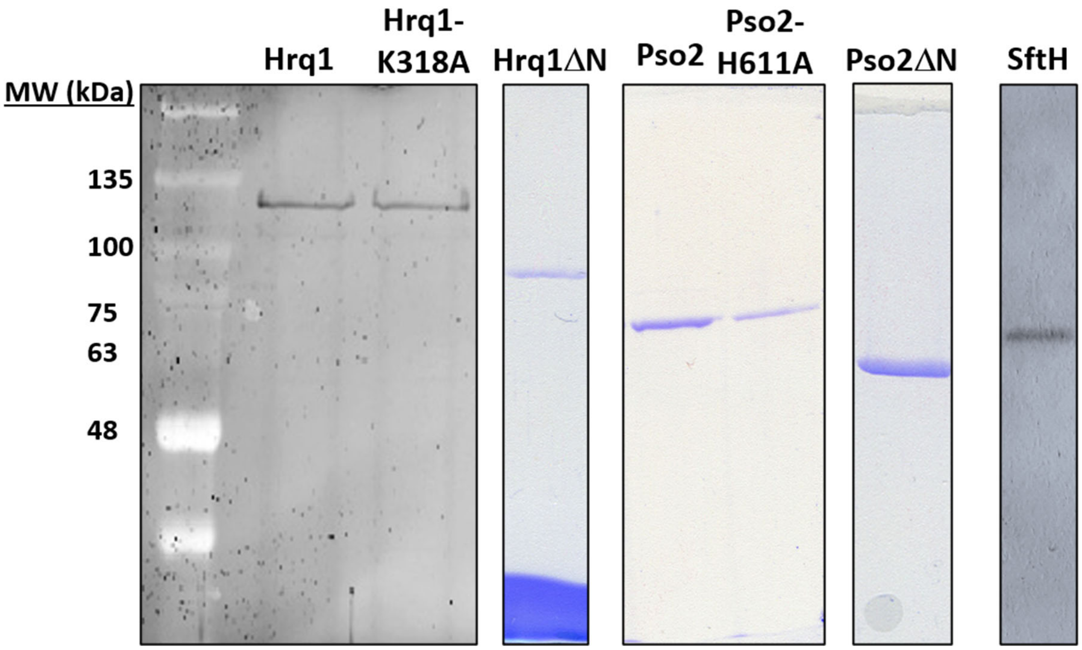


**Figure S7.** Representative SDS-PAGE images of recombinant protein preparations. The gels showing Hrq1, Hrq1-K318A, and SftH were stained with Sypro Orange, and the other gels were stained with Coomassie blue.

**Supporting tables**

**Table S1.** Oligonucleotides used in this study.

| Name | Sequence | Substrate |
|---|---|---|
| MB733 | ACCGTTGTGCAACTGAGTGGACAACGTGTCACTCACATAGCGTTC | 25-nt Random Fork, 5'-tail |
| MB734 | GAACGCTATGTGAGTGACACCAACAGGTGAGTCAACGTGTTGCCA | 25-nt Random Fork, 3'-tail |
| MB1614 | /5Phos/GACGCTGCCGAACTTAGACTTGCT | Undamaged nuclease substrate |
| MB1461 | TGAGTGAGCAAGTCTAAGTTCGGCAGCGTC | Undamaged nuclease substrate |
| MB1599 | /5Phos/GACGAC/ideoxyU/TACTGCCGAGACTTGCT | ICL-containing substrate |
| MB1600 | TGAGTGAGCAAGTCTCGGCAGTAAGTCGTC | ICL-containing substrate |
| MB1620 | /5Phos/GACGAC/ideoxyU/TACTGCCGAGACATGCTCACTCA | ICL-containing smFRET substrate, digested strand |
| MB1621 | /5Biosg/TGAGTGAGCA/iAmMC6T/GTCTCGGCAGTAAGTCGTC/3Cy3Sp/ | smFRET substrate, labelled strand |
| MB1622 | /5Phos/GACGACTTACTGCCGAGACATGCTCACTCA | Undamaged smFRET substrate, digested strand |

Abbreviations: /5Phos/, 5′ phosphate; /ideoxyU/, internal dU; ICL, inter-strand crosslink; smFRET, single-molecule Förster resonance energy transfer; /5Biosg/, 5′ biotin; and /3Cy3Sp/, 3′ Cy3 dye.

**Table S2.** *Saccharomyces cerevisiae* strains used in this study.

| Strain | Genotype | Origin |
|---|---|---|
| YHP499 | *MATa ura3-52 lys2-801_amber ade2-101_ochre trp1Δ63 his3Δ200 leu2Δ1* | (54) |
| MBY321 | *MATα ura3-52 lys2-801_amber ade2-101_ochre trp1Δ63 his3Δ200 leu2Δ1 hxt13::URA3 sgs1::his5+* | This work |
| MBY327 | *MATα ura3-52 lys2-801_amber ade2-101_ochre trp1Δ63 his3Δ200 leu2Δ1 hxt13::URA3 sgs1::his5+ hrq1::TRP1* | This work |
| MBY462 | *MATα ura3-52 lys2-801_amber ade2-101_ochre trp1Δ63 his3Δ200 leu2Δ1 hxt13::URA3 hrq1::His3MX6::Hrq1ΔN*$^{1-279}$*-NatMX* | This work |
| MBY686 | *MATa ura3-52 lys2-801_amber ade2-101_ochre trp1Δ63 his3Δ200 leu2Δ1 pso2::pso2ΔN*$^{1-94}$*-His3MX6* | This work |
| MBY745 | *MATa ura3-52 lys2-801_amber ade2-101_ochre trp1Δ63 his3Δ200 leu2Δ1 hrq1::His3MX6* | This work |
| MBY746 | *MATa ura3-52 lys2-801_amber ade2-101_ochre trp1Δ63 his3Δ200 leu2Δ1 pso2::TRP1* | This work |
| MBY747 | *MATa ura3-52 lys2-801_amber ade2-101_ochre trp1Δ63 his3Δ200 leu2Δ1 hrq1::His3MX6 pso2::TRP1* | This work |
| MBY748 | *MATa ura3-52 lys2-801_amber ade2-101_ochre trp1Δ63 his3Δ200 leu2Δ1 hrq1::hrq1-K318A-His3MX6* | This work |
| MBY749 | *MATa ura3-52 lys2-801_amber ade2-101_ochre trp1Δ63 his3Δ200 leu2Δ1 pso2::pso2-H611A-TRP1* | This work |
| MBY789 | *MATa ura3-52 lys2-801_amber ade2-101_ochre trp1Δ63 his3Δ200 leu2Δ1 pESC-URA* | This work |
| MBY790 | *MATa ura3-52 lys2-801_amber ade2-101_ochre trp1Δ63 his3Δ200 leu2Δ1 pESC-URA(PSO2)* | This work |
| MBY877 | *MATa ura3-52 lys2-801_amber ade2-101_ochre trp1Δ63 his3Δ200 leu2Δ1 pso2::TRP1 sgs1::NatMX* | This work |
| MBY878 | *MATa ura3-52 lys2-801_amber ade2-101_ochre trp1Δ63 his3Δ200 leu2Δ1 hrq1::His3MX6 pso2::TRP1 sgs1::NatMX* | This work |
